# To Play or Not to Play? Effects of Playmate Familiarity and Social Isolation on Social Play Engagement in Three Laboratory Rat Strains

**DOI:** 10.1101/2024.11.14.623692

**Authors:** Isabella C. Orsucci, Kira D. Becker, Jackson R. Ham, Jessica D.A. Lee, Samantha M. Bowden, Alexa H. Veenema

**Affiliations:** Department of Psychology, Michigan State University, East Lansing, MI 48824, USA; Department of Neuroscience, University of Lethbridge, Lethbridge, Alberta, Canada

**Author notes:** Corresponding author: Alexa H. Veenema Department of Psychology Michigan State University East Lansing MI 48824, USA. Equal contribution (Isabella C. Orsucci, Kira D. Becker). Department of Psychology and Neuroscience, University of North Carolina at Chapel Hill, Chapel Hill, NC 27514, USA.

## Abstract

Social play is a motivating and rewarding behavior displayed by juveniles of many mammalian species, including humans and rats. Social play is vital to the development of social skills. Autistic children show less social play engagement which may contribute to their impairments in social skills. There is limited knowledge about what external conditions may positively or negatively influence social play engagement in humans or other animals. Therefore, we determined how two common external conditions, playmate familiarity and social isolation, modulate social play levels and social play defense tactics in juveniles of three common laboratory rat strains: Long-Evans, Sprague-Dawley, and Wistar. Males and females were socially isolated for either 2h or 48h prior to social play testing and were then exposed to either a familiar (cage mate) or novel playmate, creating four testing conditions: 2h-Familiar, 48h-Familiar, 2h-Novel, and 48h-Novel. Both playmate familiarity and social isolation length influenced social play behavior levels and tactics in juvenile rats, but did so differently for each of the three rat strains. Long-Evans played most with a familiar playmate, irrespective of time isolated, Sprague-Dawley played most in the 48h-Familiar condition, and Wistar played the least in the 2h-Familiar condition, but Wistar played more with a novel playmate than Long-Evans and Sprague-Dawley. Analysis of social play tactics by the playmates in response to nape attacks by the experimental rats revealed strain differences with novel playmates. Here, Sprague-Dawley and Wistar defended more nape attacks than Long-Evans. Sprague-Dawley evaded these attacks, thereby shortening body contact. In contrast, Wistar turned to face their playmate attacker and showed more complete rotations, thereby extending body contact and wrestling longer. Role reversals, which increase social play reciprocity and reflect the quality of social play, were higher in Long-Evans and Sprague-Dawley with familiar playmates. Role reversals decreased for Sprague-Dawley but increased for Wistar after 48h isolation. The effects of playmate familiarity or social isolation length on social play levels and tactics were similar across sex for all three strains. In conclusion, we showed that two common external factors (playmate familiarity and social isolation length) that largely vary across social play studies have a major impact on the level and quality of social play in the three rat strains. Strain differences indicate higher level and quality of social play with familiar playmates in Long-Evans, with familiar playmates after short isolation in Sprague-Dawley, and with novel playmates after longer isolation for Wistar. Future research could determine whether strain differences in neuronal mechanisms underlie these condition-induced variations in social play engagement. Our findings are also informative in suggesting that external conditions like playmate familiarity and social isolation length could influence social play levels and social play quality in typical and atypical children.

## INTRODUCTION

Social play, also known as rough-and-tumble play or play-fighting, is the most characteristic social behavior in young mammals (Manduca et al., 2014; Schiavi et al., 2020), with juvenile animals spending upwards of 20% of their time participating in play (Pellis and Pellis, 2013). Social play is a highly motivating and rewarding behavior (Calcagnetti and Schechter, 1992; Pellis and Pellis, 1987; Trezza et al., 2010; Vanderschuren et al., 2016). The pleasures of play have been long recognized by scientists; Charles Darwin wrote that ‘Happiness is never better exhibited than by young animals, such as puppies, kittens and lambs, when playing together, like our own children’ (Darwin et al., 1871). Nonetheless, social play does much more than bring pleasure to young mammals. In humans, non-human primates, and rats, social play is important to the development of emotional, social, physical, and cognitive skills (Barnett, 1990; Ginsburg et al., 2007; Ljubetic et al., 2020; Nijhof et al., 2018; Pellis et al., 2023, 2014; Spinka et al., 2001; Vanderschuren et al., 1997). Social play also promotes executive function, contributes to the development of a prosocial brain, and helps to develop flexible problem solving skills (Barnett, 1990; Ham et al., 2024a; Panksepp, 2007, 1998; Pellis et al., 2017; Spinka et al., 2001; Yogman et al., 2018).

Social play deficits are a core symptom of neurodevelopmental disorders, such as autism spectrum disorder (ASD), early-onset schizophrenia, and attention-deficit/hyperactivity disorder (Alessandri, 1992; Helgeland and Torgersen, 2005; Jones, 1994; Jordan, 2003; Moller and Husby, 2000). For example, ASD children show less involvement in social play with their peers compared to typically developing children (Alessandri, 1992; Moller and Husby, 2000; Panksepp, 1981; Supekar et al., 2018). ASD children also report that they find social interactions less pleasant than non-social interactions, such as bike-riding or eating (Supekar et al., 2018). Furthermore, schizophrenic children show no attempt to establish social play with other children (Potter, 1933), and do not show interest in playing with school mates (Potter, 1933). Moreover, the preference for playing alone versus with others in children ages four to six predicted schizophrenia (Bender, 1947; Jones et al., 1995). It has been suggested that limited social play engagement may reduce the development of social skills and thus the ability of ASD and schizophrenic individuals to appropriately navigate social and emotional situations later in life (Jordan, 2003). This indicates the need to enhance our understanding of what conditions influence social play engagement. The play environment is an external condition that can influence social play engagement in ASD children. For example, enclosed/confined play environments increased peer interaction and play between ASD children (Black et al., 1975), as did playgrounds designed to have high spatial density and a structured layout (Yuill et al., 2007). Another external condition is the quality of the play partner. Here, social interactions, including social play, increase when socially competent peers are included as the play partner in ASD children (Wolfberg and Schuler, 1993). But overall, there is an incomplete understanding of what specific external conditions may positively or negatively influence engagement in social play in human children or other young animals.

The vast majority of our current knowledge of social play behavior stems from laboratory rat studies (Siviy and Panksepp, 2011; Trezza et al., 2010; Vanderschuren et al., 2016). Social play bouts in rats typically begin with a nape attack, in which a rat touches the nape of the neck of a conspecific with its snout (Panksepp and Beatty, 1980; Pellis and Pellis, 1987; Poole and Fish, 1975; Vanderschuren et al., 1997). This is considered the most important parameter of play initiation (Vanderschuren et al., 2016). To protect the nape, the recipient of a playful attack may engage in various defensive tactics. The recipient may evade an attack, running away from the attacker. Alternatively, the recipient may turn to face its attacker. After turning to face, this frequently results in the recipient rolling onto its dorsal surface so that the nape is pressed against the floor/ground and inaccessible to the attacker. In other cases, the recipient may turn and face and meet its attacker in an upright position, where the animals appear to be boxing with one another. Regardless of the defense tactic employed, the recipient rat frequently attempts to gain access to the attacker rats’ neck area, so that play is prolonged. In turn, this maintains a level of reciprocity where the attacking rat and defending rat switch roles.

The dyad test is the most widely used experimental paradigm for the analysis of social play in rats, and typically consists of two rats joined together following a period of social isolation to increase their motivation to engage in play (Pellis et al., 2022). But even within this dyadic social play paradigm there exist many different versions in which several conditions can be modified, including the type of playmate (e.g., familiar versus novel), test enclosure (e.g., familiar versus novel), time of day at testing (dark phase versus light phase), illumination levels (bright light versus dim light versus red light versus complete darkness). Modifying one or more of these conditions may affect both the amount and style of social play behaviors (Ham et al., 2024b; Himmler et al., 2014a; Panksepp, 1981; Reinhart et al., 2006; Siviy et al., 2017).

Social play levels and social play tactics may also depend on the strain of rats. Indeed, the three standard laboratory rat strains Long-Evans, Sprague-Dawley, and Wistar differ in their social play behaviors (Himmler et al., 2013a, 2014a, 2014b). For example, Sprague-Dawley rats play more than Wistar rats when exposed to a familiar conspecific in a novel environment after 24h isolation, and being tested in complete darkness (Himmler et al., 2014b). Moreover, Sprague-Dawley rats showed higher engagement in social play than Long-Evans rats when exposed to a novel conspecific in a novel environment without prior isolation, and being tested in the light phase (Ku et al., 2016). However, Wistar and Long-Evans rats played more than Sprague-Dawley rats when exposed in their home cage to an unfamiliar sibling without prior isolation, and being tested in the light phase (Northcutt and Nwankwo, 2018). Regardless of strain, the degree of competition and cooperation, as measured by the likelihood of a role reversal occurring after a playful attack, seems similar among these three rat strains (Himmler et al., 2016, 2014b). However, other social play tactics differ among Long-Evans, Sprague-Dawley and Wistar (Himmler et al., 2013a, 2016). For example, Sprague-Dawley rats are more likely to evade a playful attack than both Long-Evans and Wistar rats (Himmler et al., 2016, 2014b). Conversely, Long-Evans and Wistar rats typically defend themselves by rolling to supine, resulting in the attacker pinning the defender (Himmler et al., 2016, 2014b). Together, these findings suggest that playmate familiarity and social isolation are both important factors influencing social play levels and tactics differently across these three rat strains.

Therefore, in the current study, we determined how two external conditions would modulate social play levels and social play tactics in Long-Evans, Sprague-Dawley, and Wistar rats. The first external condition manipulated was playmate familiarity, in which experimental rats were exposed to either a familiar (cage mate) or novel playmate during the social play tests. The second external condition manipulated was length of social isolation. Socially isolating rats for various lengths has resulted in differences in social play levels. For example, Long-Evans and Wistar rats show more social play after a 24h social isolation versus no isolation (Long-Evans) or versus 1h and 3.5h isolation (Wistar) (Panksepp and Beatty, 1980; Vanderschuren et al., 1995a). In the current study, rats were isolated for either 2h or 48h prior to the social play test.

## METHODS

### Subjects

Three-week-old male and female Long-Evans rats were obtained from Envigo (Indianapolis, IN; n=5 per sex) or were bred and raised in house (n=13 males, n=14 females). Three-week-old male and female Sprague-Dawley rats were obtained from Charles River Laboratories (Raleigh, NC; n=5 males, 5 females) or were bred and raised in house (n=9 males, 20 females). Three-week-old male and female Wistar rats were obtained from Charles River Laboratories (Raleigh, NC; n=20 males, n=20 females). Rats were maintained under standard laboratory conditions (12 hr light/dark cycle, lights off at 14:00 hr, food and water available *ad libitum*). Rats were housed in single-sex groups of three in standard Allentown cages (58.8 cm x 19.4 cm x 39.5 cm) with woodchip bedding unless otherwise mentioned. The experiments were conducted in accordance with the National Institute of Health *Guidelines for Care and Use of Laboratory Animals* and approved by Michigan State University Institutional Animal Care and Use Committee.

### Social Play Testing

Social play was assessed in postnatal day 29-34 (juvenile) rats because social play is at its peak at this age (Panksepp, 1981; Paul et al., 2014; Pellis and Pellis, 1987; Thor and Holloway, 1984). Social play tests started at the beginning of the dark phase under red-light illumination because rats are nocturnal and most active in the dark phase (Hawkins and Golledge, 2018; Himmler et al., 2013a). All testing was completed in the experimental rat’s home cage to abolish the suppression of play induced by unfamiliarity to the test cage (Vanderschuren et al., 1995b). When testing began, the experimental rat’s home cage was removed from the cage rack, placed on the testing table and its lid was replaced with a Plexiglass lid to allow recording of the tests from above using a tripod and video camera. A sex-, age-, and strain-matched playmate was placed into the experimental rat’s home cage. The experimental subject was allowed to freely interact with the stimulus subject for 10 min. Food and water were not available during the 10 min testing sessions but were immediately returned when each session was complete. Rats assigned playmates were striped on their back and sides with a Sharpie marker 1h prior to social play testing to distinguish between the experimental and playmates during later video analysis.

### Behavioral Scoring

The 10-min social play tests were videotaped, and the behavior of the experimental rats was measured by a researcher blind to the test conditions using SolomonCoder software (https://solomon.andraspeter.com/). The following social play behaviors were scored for the experimental rats according to Veenema and Neumann (2009): duration of social play (the total amount of time spent in playful social interactions including nape attacks, pinning, and supine poses), numbers of nape attacks (nose attacks or nose contacts towards the nape of the neck of the intruder), number of pins (the resident holds the intruder on its back in a supine position), and number of supine poses (the resident is pinned by the intruder). Other behaviors scored were duration of social investigation (sniffing the anogenital and head/neck regions), duration of allogrooming (the experimental subject is grooming the stimulus subject), and duration of non-social cage exploration (the experimental subject is walking, rearing, sitting, or engaging in other neutral behaviors). Finally, the duration of total social behaviors (social play duration, social investigation duration, allogroom duration) was calculated. Results of these other behaviors and total social behaviors appear in the supplementary materials and are not further discussed in the main document.

Additionally, social play tactics used to defend against a nape attack of the experimental rat were scored for the experimental rat’s playmate, using a combination of normal speed and frame-by-frame analysis following the protocol described by (Pellis et al., 2022). After a nape attack is launched by the experimental rat, the playmate can employ different defensive tactics (Figure 1). If the playmate defends itself, it can either engage in an evasive maneuver (i.e., runs away) or it can turn to face its attacker (i.e., facing defense) (Himmler et al., 2013b). If the playmate turns to face, it can do so by either rotating along the vertical or horizontal axis of the body. If the playmate rotates along the vertical axis, the pair ends in an upright, standing defense position, where they appear as if they are boxing. If the playmate rotates along the horizontal axis, the pair ends in one of two pin configurations: a partial rotation, where the playmate’s body is rotated ∼45°, or a complete rotation, where the playmate is completely supine and has rotated ∼90°. Both pin measures indicate the playmate’s motivation to sustain in close bodily contact with the attacker rat (Stark et al., 2021). Finally, rats engage in role reversals or turn taking, and the degree to which role reversals are engaged can serve as a measure of reciprocity and how motivated the playmate is to continue the ongoing playful encounter (Pellis et al., 2024). Using these behavioral definitions, for each playmate, the proportion of nape attacks that were defended were calculated. Of the defended attacks, the proportion that led to a facing defense was calculated, along with the proportion that were evaded, leading to proportions of partial and complete rotations, and mutual uprights that were calculated. Lastly, the proportion of playful attacks that were the result of a role reversal were calculated. See Ham and Pellis (2024) for a detailed description of the calculations.

**Figure 1.**
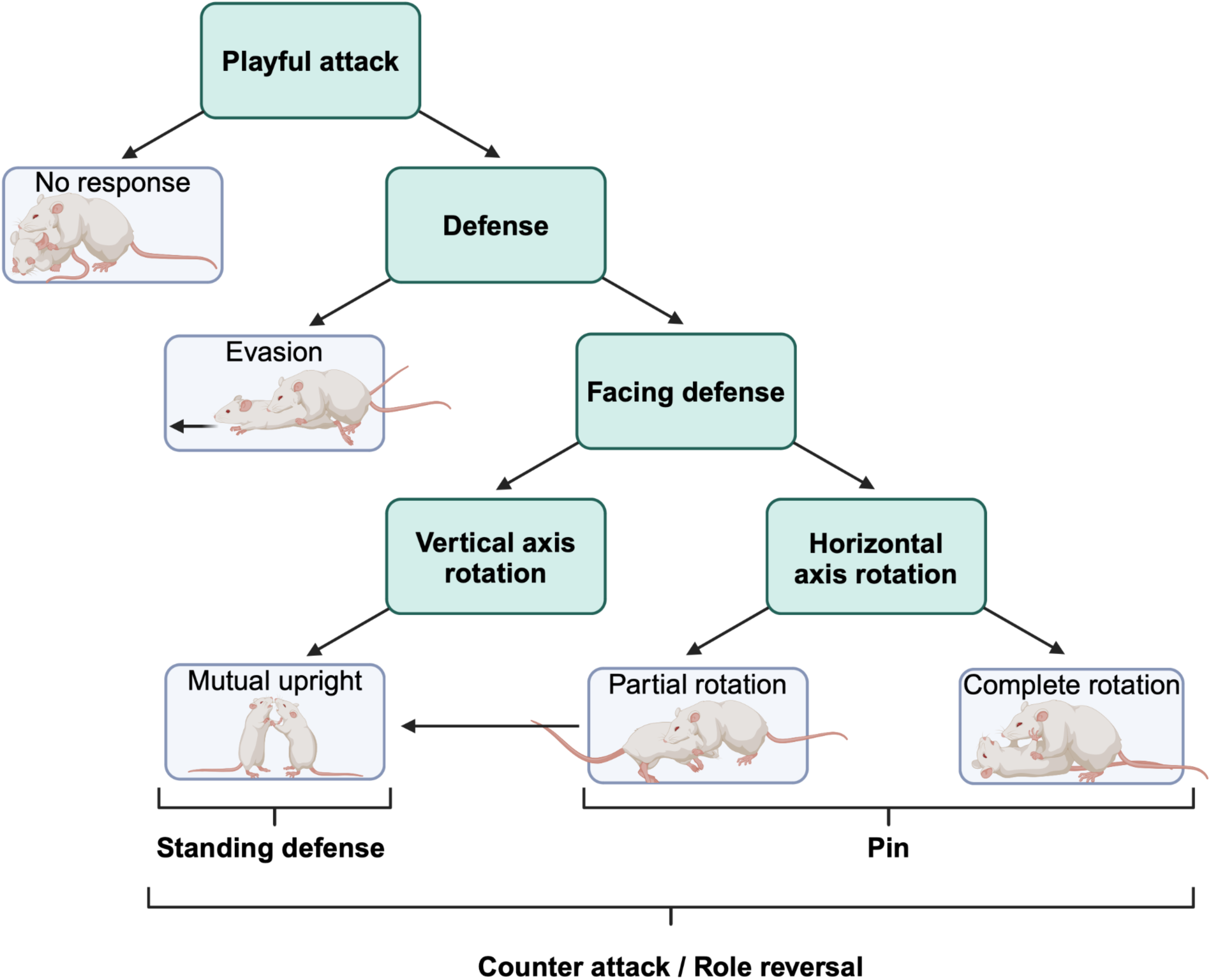
Decision tree showing the diverse social play defense tactics of the experimental rat’s playmate in response to a nape attack by the experimental rat. When attacked, a rat can either defend itself (defense) or ignore the playful attack (no response). If the attacked rat defends itself, it can either show evasion (i.e., run away) or engage in a facing defense. If it turns to face the attacker rat, it either rotates along the vertical body axis or horizontal body axis. Vertical axis rotations end in a mutual upright or standing defense. In contrast, horizontal axis rotations result in either a partial or complete rotation, both of which are commonly referred to as a pin. Only attacks that are defended can result in a role reversal. For an in-depth read of the microstructure of rat play, see Pellis et al. (2022). Adapted from Stark (2021).

### Experimental Conditions

Experimental rats were randomly assigned to one of four experimental conditions in which length of social isolation (2h or 48h) and familiarity of playmate (cage mate or novel playmate) were fixed factors, resulting in four experimental groups: (1) 2h isolation + familiar playmate (2h-Familiar), (2) 48h isolation + familiar playmate (48h-Familiar), (3) 2h isolation + novel playmate (2h-Novel), and (4) 48h isolation + novel playmate (48h-Novel). For the isolation periods, the experimental rat stayed in its home cage while the two cagemates were removed and placed together in a new cage. 2h or 48h later, the 10-min social play tests were conducted by placing either a cagemate (familiar playmate) or a sex- and age-matched novel playmate in the experimental rat’s home cage. All playmates (cagemates and novel playmates) were group housed prior to being placed into the experimental’s home cage for social play testing. After testing was completed, the two cagemates were rehoused with the experimental rat in the experimental rat’s home cage. See Figure 2 for experimental design and Table 1 for number of rats per experimental condition.

**Figure 2.**
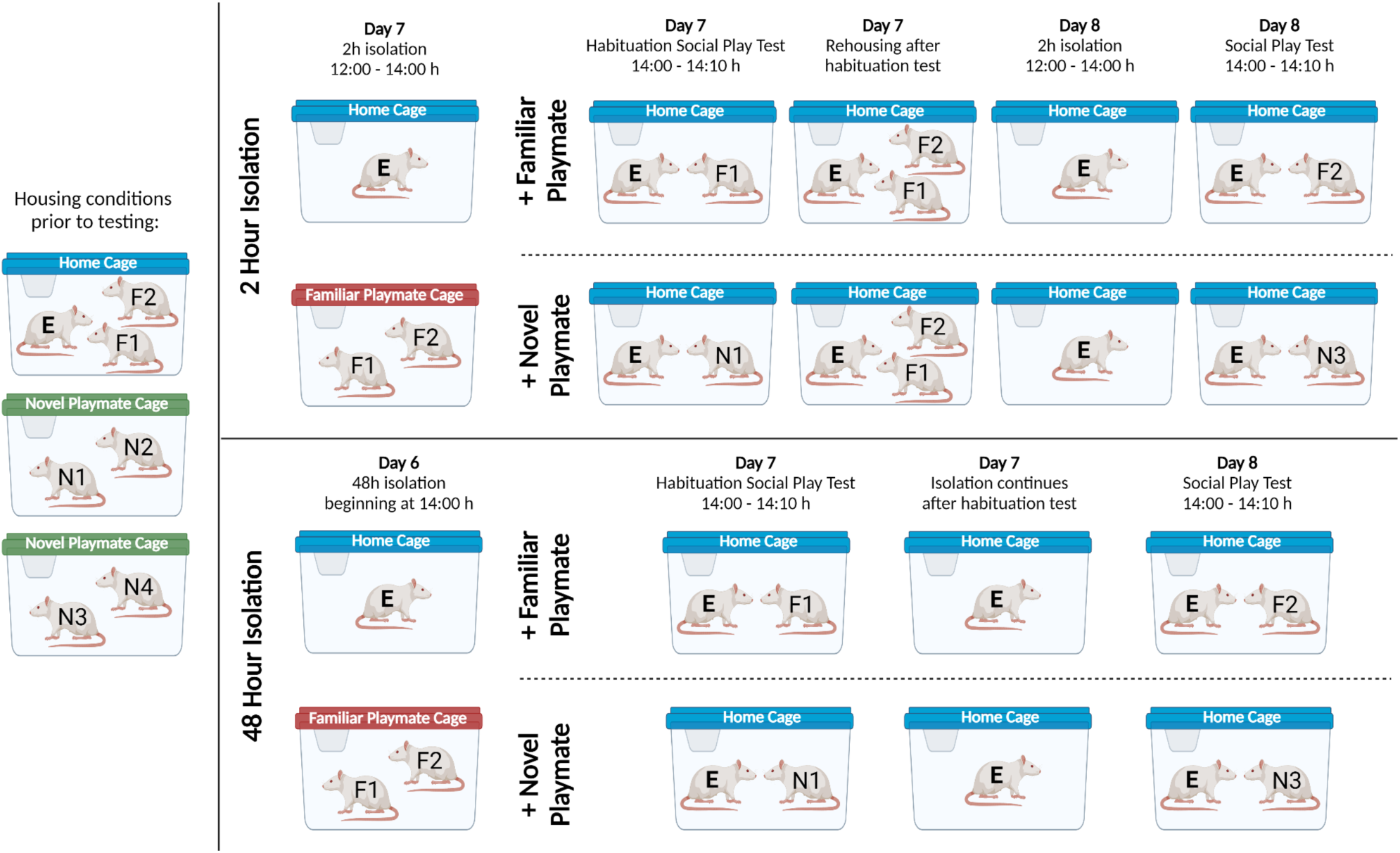
Experimental design to determine the effects of playmate familiarity (familiar or novel) and social isolation (2h or 48h) on social play engagement in juvenile male and female Long-Evans, Sprague-Dawley and Wistar rats. On Day 1, rats were randomly assigned as experimental rat (E), familiar playmate (F1, F2), or novel playmate (N1-4). During social isolation, the experimental rat remained in its home cage (indicated with a blue lid) while the two cagemates (F1 and F2) were placed in a new cage (‘familiar playmate cage’, indicated with a red lid). Novel playmates were housed in pairs (‘novel playmate cages’, indicated with a green lid). Social play testing occurred in the home cage of the experimental rat at the start of the dark phase (14:00 h). After a 2h or 48h isolation period, experimental rats were exposed to a 10- min Habituation Social Play Test on Day 7 with either a familiar (F1) or novel (N1 or N2) playmate. After this habituation test, the familiar playmates were rehoused with the experimental rats in the 2h Isolation group while the experimental rats in the 48h Isolation group remained singly housed. On Day 8, experimental rats underwent the 10-min Social Play Test by exposing them to the other familiar playmate (F2) or to a novel playmate from the other novel playmate cage (N3 or N4). All familiar and novel playmates remained socially housed during the experiment to prevent any isolation effects on the stimulus rats. Image created with BioRender.com

**Table 1.** Number of rats per strain, sex, and experimental condition.

|  | 2h-Familiar | 48h-Familiar | 2h-Novel | 48h-Novel |
| --- | --- | --- | --- | --- |
| Long-Evans<br>(n=37) | Male: 4<br>Female: 6 | Male: 5, 4<br>Female: 5, 4 | Male: 5<br>Female: 5 | Male: 5<br>Female: 4 |
| Sprague-Dawley<br>(n=39) | Male: 3<br>Female: 7 | Male: 5, 2<br>Female: 5, 5 | Male: 5<br>Female: 5 | Male: 4<br>Female: 8 |
| Wistar | Male: 10 | Male: 10 | Male: 10 | Male: 10 |
| (n=40) | Female: **10** | Female: *10* | Female: *10* | Female: **10** |

Not every rat was tested in every condition. Bold or italic font represents the same rats used in two conditions. Normal font represents rats tested in one condition.

### Statistical Analysis

Statistical analysis was done using IBM SPSS. A three-factor design analysis of variance (ANOVA) was conducted to determine the effects of Playmate (familiar versus novel), Isolation (2h versus 48h), and Sex on social play expression and social play tactics per strain. A one factor design ANOVA was conducted to determine the effects of Strain (Long-Evans, Sprague-Dawley, Wistar) on social play expression and social play tactics for each of the four experimental conditions (2h-Familiar, 48h-Familiar, 2h-Novel, 48h-Novel). When significant interactions or strain differences were found, Bonferroni post hoc tests were conducted to clarify the effects. Significance was set at p < 0.05. Partial eta squared (ƞ_p_^2^) were manually computed when significant main effects or interactions were found.

## RESULTS

### 1. Experimental Condition comparisons per rat strain for social play behavior

#### Long-Evans

There was a significant main effect of Playmate on the duration of social play, number of nape attacks, and number of pins (See Table 2 for statistical details), with more social play, nape attacks and pins with exposure to a familiar versus novel playmate (Fig 3A-C).

**Fig. 3.**
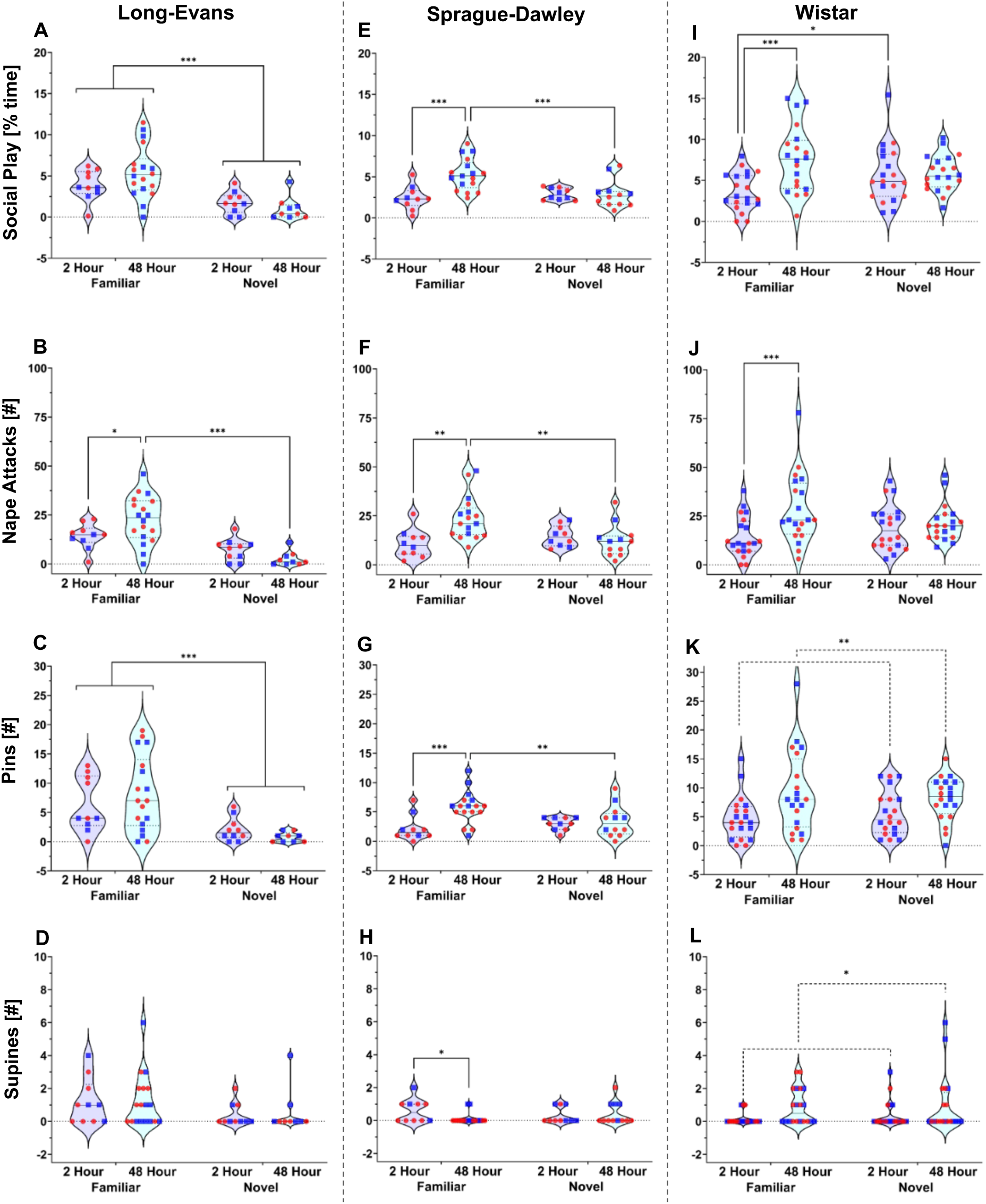
Experimental condition comparisons for social play behaviors per rat strain - Long-Evans showed most social play with familiar playmates, Sprague-Dawley show most social play with familiar playmates after 48h isolation, and Wistar showed the least social play with familiar playmates after 2h isolation. **A-D:** Long-Evans played more with a familiar than a novel playmate, as shown by longer duration of social play under both 2h and 48h isolation **(A)**, more nape attacks under 48-h isolation **(B)**, and more pins under both 2h and 48h isolation **(C)** while no effects were seen for number of supines **(D)**. **E-G:** Sprague-Dawley played more with a familiar playmate after 48h isolation than with a familiar playmate after 2h isolation and a novel playmate after 48h isolation, as shown by longer duration of social play **(E)**, more nape attacks **(F),** and more pins **(G)**. Sprague-Dawley had more supines with a familiar playmate after 2h versus 48h isolation (**H)**. **I-J**: Wistar played more with a familiar playmate after 48h versus 2h isolation, as shown by longer duration of social play **(I)** and more nape attacks **(J)**. Wistar played more with a novel than a familiar playmate after 2h isolation **(I)**. **K-L**: Wistar had more pins **(K)** and supines **(L)** after 48h versus 2h isolation. Males are represented by blue squares, females are represented by red circles; Three-way ANOVA analyses per strain to assess effects of Playmate, Isolation, and Sex, followed by Bonferroni post hoc tests to assess interaction effects; * *p* < 0.05, ** *p* ≤ 0.01, *** *p* ≤ 0.001.

**Table 2.** Experimental Condition comparisons for social play behaviors in experimental Long-Evans rats. Three-way ANOVA statistics and partial eta squared (η2) effect sizes for the effects of Playmate (Familiar, Novel), Isolation (2h, 48h), and Sex on behavior of experimental male and female juvenile Long-Evans rats in the social play test. Significant effects and their corresponding behaviors are indicated in **bold.**

|  | Playmate | Isolation | Playmate x Isolation | Sex |
| --- | --- | --- | --- | --- |
| **Social play**<br>**(% Time)** | **$F_{(1,39)} = 20.4$,**<br>**$p < 0.001$**<br>**$\eta_p^2 = 0.34$** | $F_{(1,39)} = 0.42$,<br>$p = 0.520$ | $F_{(1,39)} = 3.03$,<br>$p = 0.090$ | $F_{(1,39)} = 0.90$,<br>$p = 0.350$ |
| **Nape**<br>**attacks**<br>**(#)** | **$F_{(1,39)} = 24.3$,**<br>**$p < 0.001$**<br>**$\eta_p^2 = 0.38$** | $F_{(1,39)} = 0.56$,<br>$p = 0.457$ | **$F_{(1,39)} = 6.44$,**<br>**$p = 0.015$**<br>**$\eta_p^2 = 0.14$** | $F_{(1,39)} = 1.23$,<br>$p = 0.274$ |
| **Pins**<br>**(#)** | **$F_{(1,39)} = 15.2$,**<br>**$p < 0.001$**<br>**$\eta_p^2 = 0.28$** | $F_{(1,39)} = 0.15$,<br>$p = 0.701$ | $F_{(1,39)} = 1.73$,<br>$p = 0.197$ | $F_{(1,39)} = 2.17$,<br>$p = 0.148$ |
| Supine<br>poses<br>(#) | $F_{(1,39)} = 3.40$,<br>$p = 0.073$ | $F_{(1,39)} = 0.01$,<br>$p = 0.933$ | $F_{(1,39)} = 0.02$,<br>$p = 0.882$ | $F_{(1,39)} = 0.42$,<br>$p = 0.523$ |

There was a significant Playmate x Isolation effect on the number of nape attacks (Table 2). Bonferroni post hoc tests indicate that Long-Evans in the 48h-Familiar condition showed more nape attacks compared to the 2h-Familiar and 48h-Novel conditions (Fig. 3B; Table 3).

**Table 3.**
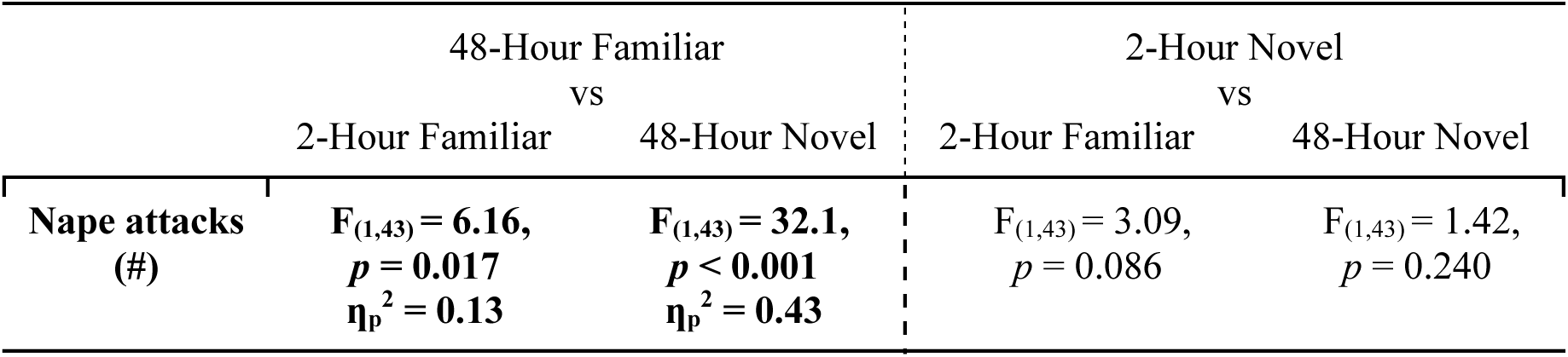
Posthoc tests for experimental condition comparisons for social play behaviors in experimental Long-Evans rats. Bonferroni post hoc statistics and partial eta squared (η2) effect sizes for the interaction effects of Playmate (Familiar, Novel) and Isolation (2h, 48h) on behaviors of experimental male and female juvenile Long-Evans rats in the social play test. Significant effects and their corresponding behaviors are indicated in **bold**.

There were no significant main effects of Sex or interaction effects with Sex on any social play behavior analyzed (Table 2).

#### Sprague-Dawley

There was a significant main effect of Isolation on the duration of social play and the number of pins (Table 4), with more social play and pins after 48h isolation versus 2h isolation (Fig. 3E,3G).

**Table 4.**
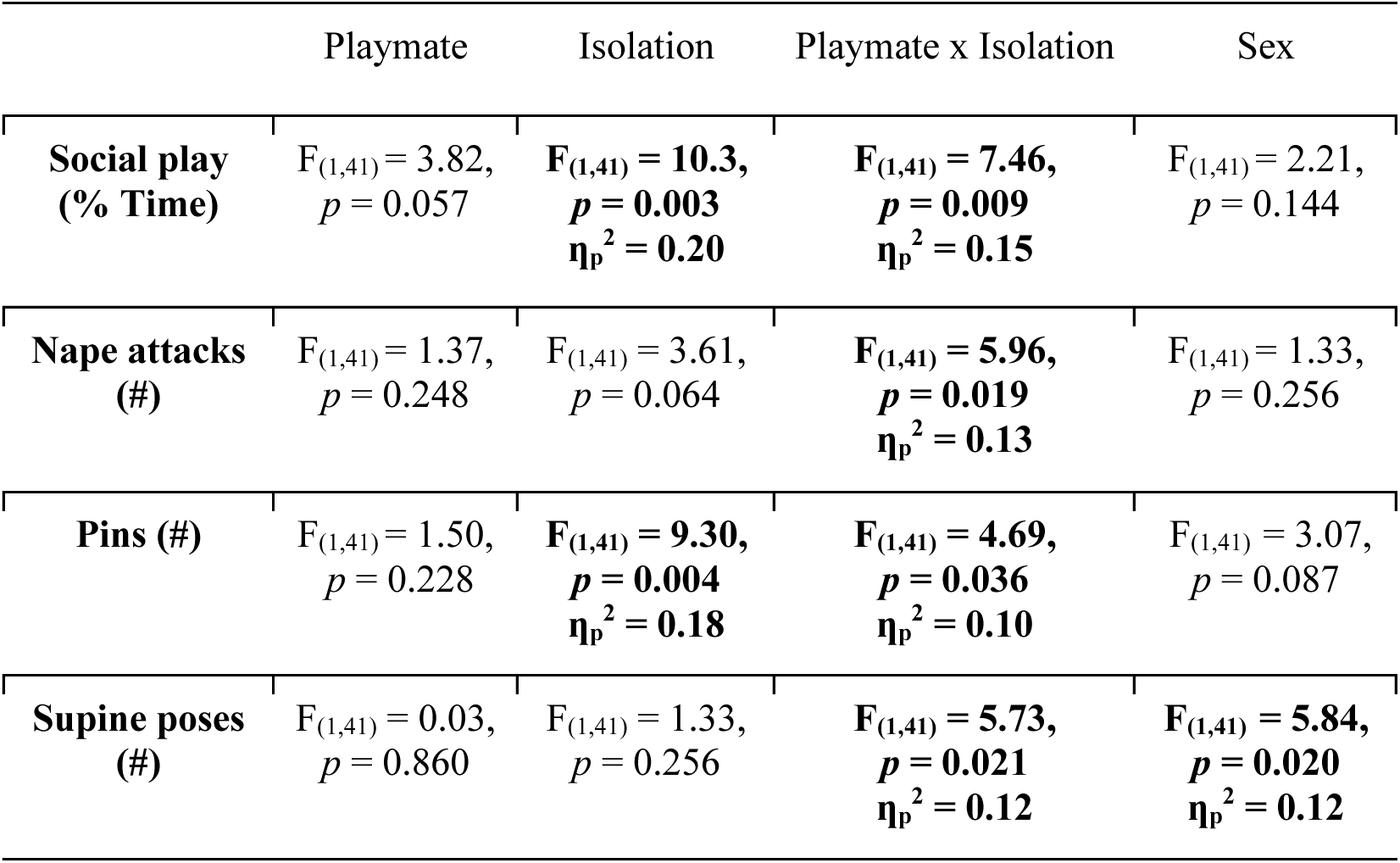
Experimental Condition comparisons for social play behaviors in experimental Sprague-Dawley rats. Three-way ANOVA statistics and partial eta squared (η2) effect sizes for the effects of Playmate (Familiar, Novel), Isolation (2h, 48h), and Sex on behavior of experimental male and female juvenile Sprague-Dawley rats in the social play test. Significant effects and their corresponding behaviors are indicated in **bold**.

There was a significant Playmate x Isolation effect on the duration of social play, number of nape attacks, number of pins, and number of supine poses (Table 4). Bonferroni post hoc tests indicate that Sprague-Dawley in the 48h-Familiar condition showed more social play, nape attacks, and pins compared to the 2h-Familiar and the 48h-Novel conditions (Fig. 3E-G; Table 5) but fewer supine poses compared to the 2h-Familiar condition (Fig. 3H; Table 5).

**Table 5.**
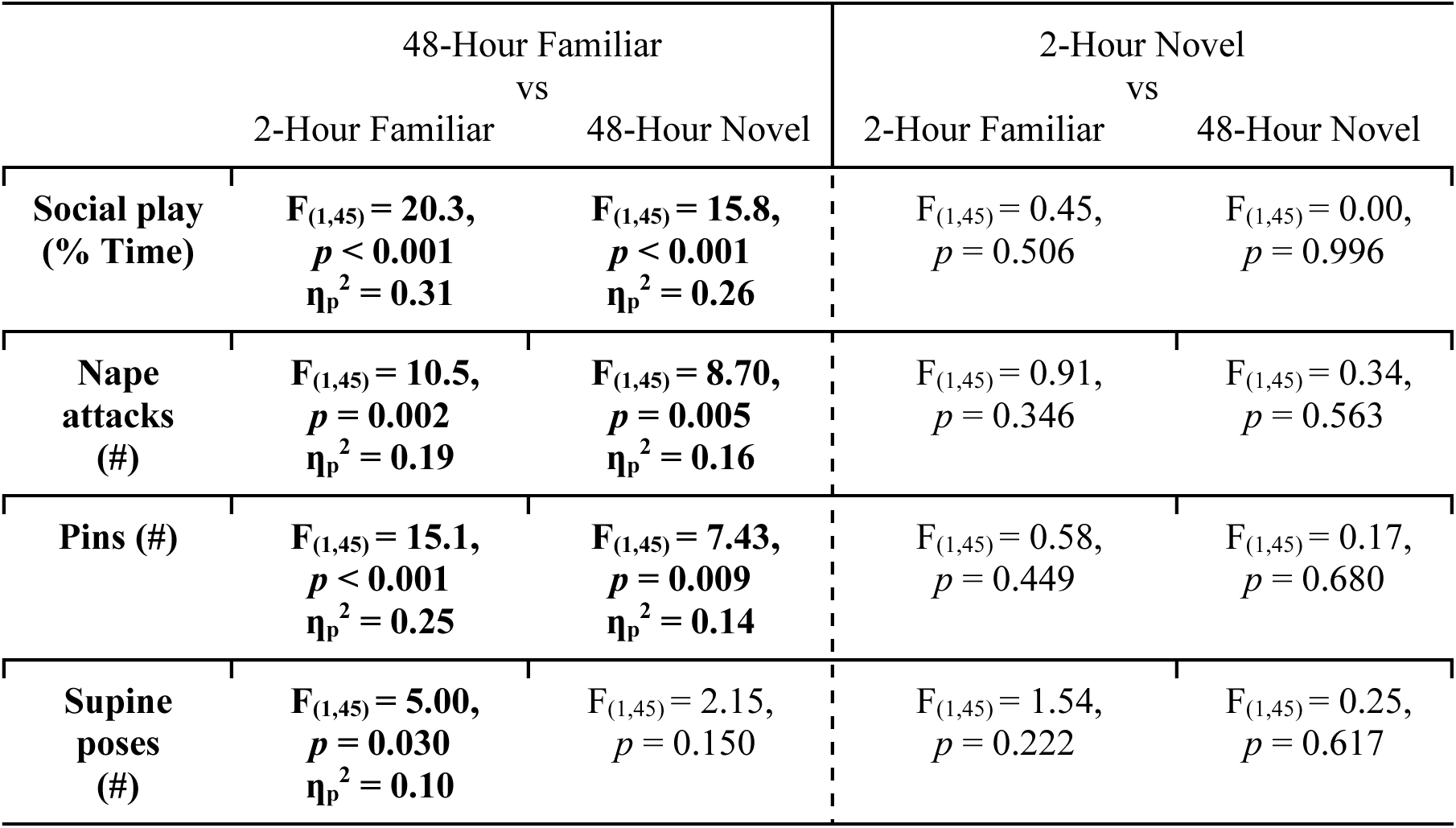
Posthoc tests for experimental condition comparisons for social play behaviors in experimental Sprague-Dawley rats. Bonferroni post hoc statistics and partial eta squared (η2) effect sizes for the interaction effects of Playmate (Familiar, Novel) and Isolation (2h, 48h) on behavior of experimental male and female juvenile Sprague-Dawley rats in the social play test. Significant effects and their corresponding behaviors are indicated in **bold**.

There was a significant main effect of Sex on the number of supine poses (Table 4), with more supine poses in males versus females.

#### Wistar

There was a significant main effect of Isolation on the duration of social play, number of nape attacks, pins, and supine poses (Table 6), with more social play, nape attacks, pins, and supine poses after 48h isolation versus 2h isolation (Fig. 3I-L).

**Table 6.** Experimental Condition comparisons for social play behaviors in experimental Wistar rats. Three-way ANOVA statistics and partial eta squared (η2) effect sizes for the effects of Playmate (Familiar, Novel), Isolation (2h, 48h), and sex on behavior of experimental male and female juvenile Wistar rats in the social play test. Significant effects are indicated in **bold**.

|  | Playmate | Isolation | Playmate x Isolation | Sex |
| --- | --- | --- | --- | --- |
| **Social play**<br>**(% Time)** | $F_{(1,72)} = 0.06,$<br>$p = 0.811$ | $F_{(1,72)} = \mathbf{9.43},$<br>$p = \mathbf{0.003}$<br>$\eta_p^2 = \mathbf{0.12}$ | $F_{(1,72)} = \mathbf{7.73},$<br>$p = \mathbf{0.007}$<br>$\eta_p^2 = \mathbf{0.10}$ | $F_{(1,72)} = \mathbf{5.93},$<br>$p = \mathbf{0.017}$<br>$\eta_p^2 = \mathbf{0.08}$ |
| **Nape attacks**<br>**(#)** | $F_{(1,72)} = 0.16,$<br>$p = 0.695$ | $F_{(1,72)} = \mathbf{8.43},$<br>$p = \mathbf{0.005}$<br>$\eta_p^2 = \mathbf{0.10}$ | $F_{(1,72)} = \mathbf{4.55},$<br>$p = \mathbf{0.036}$<br>$\eta_p^2 = \mathbf{0.06}$ | $F_{(1,72)} = \mathbf{5.94},$<br>$p = \mathbf{0.017}$<br>$\eta_p^2 = \mathbf{0.08}$ |
| **Pins (#)** | $F_{(1,72)} = 0.01,$<br>$p = 0.927$ | $F_{(1,72)} = \mathbf{10.2},$<br>$p = \mathbf{0.002}$<br>$\eta_p^2 = \mathbf{0.12}$ | $F_{(1,72)} = 0.85,$<br>$p = 0.359$ | $F_{(1,72)} = 2.33,$<br>$p = 0.132$ |
| **Supine poses**<br>**(#)** | $F_{(1,72)} = 0.36,$<br>$p = 0.550$ | $F_{(1,72)} = \mathbf{6.80},$<br>$p = \mathbf{0.011}$<br>$\eta_p^2 = \mathbf{0.09}$ | $F_{(1,72)} = 0.16,$<br>$p = 0.690$ | $F_{(1,72)} = 0.04,$<br>$p = 0.842$ |

There was a significant Playmate x Isolation effect on the duration of social play and the number of nape attacks (Table 6). Bonferroni post hoc tests indicate that Wistar in the 2h-Familiar condition showed less social play compared to the 2h-Novel and 48h-Familiar conditions and fewer nape attacks compared to the 48h-Familiar condition (Fig. 3I-J; Table 7).

**Table 7.**
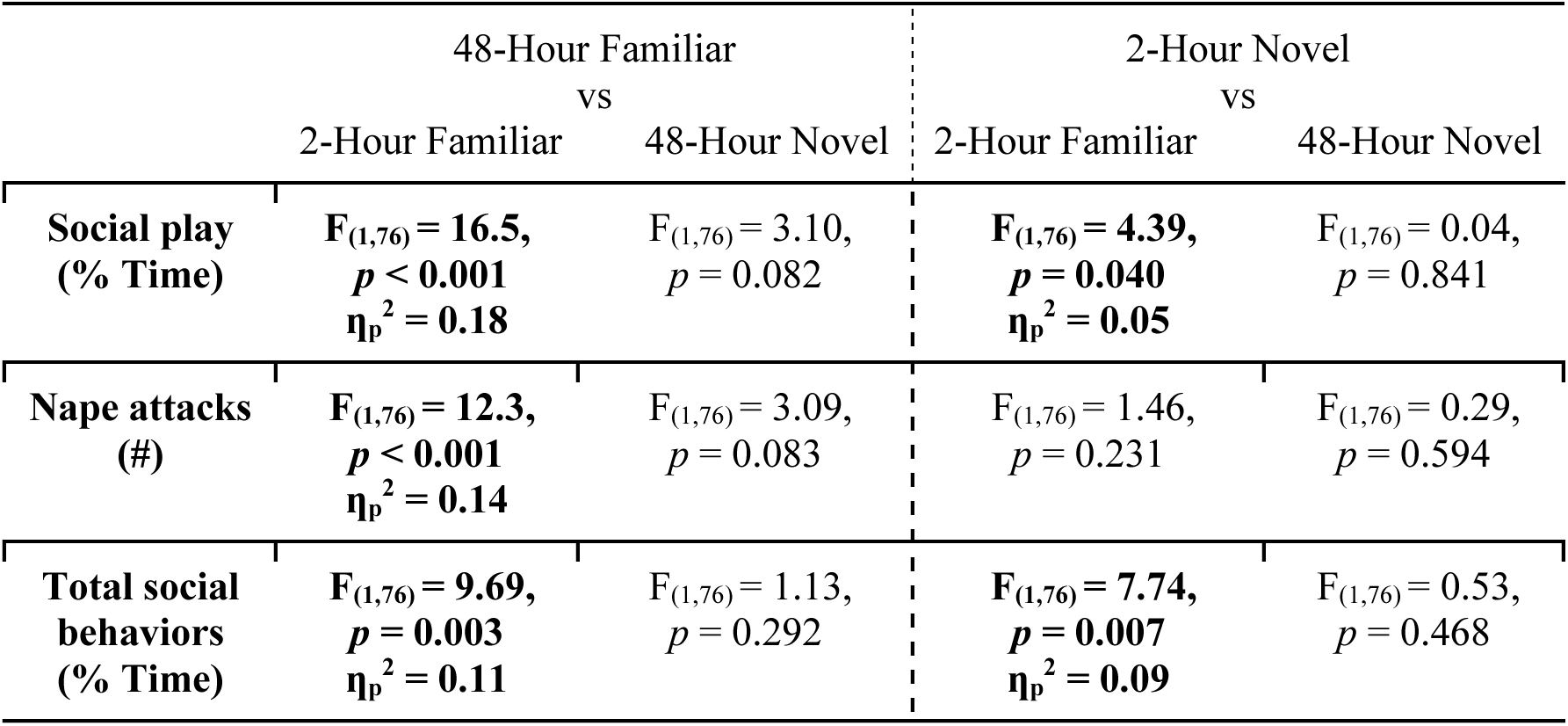
Posthoc tests for experimental condition comparisons for social play behaviors in experimental Wistar rats. Bonferroni post hoc statistics and partial eta squared (η2) effect sizes for the interaction effects of Playmate (Familiar, Novel) and Isolation (2h, 48h) on behavior of experimental male and female juvenile Wistar rats in the social play test. Significant effects are indicated in **bold**.

There was a significant main effect of Sex on the duration of social play and the number of nape attacks (Table 6), with more social play and more nape attacks in males versus females.

### 2. Experimental Condition comparisons per rat strain for social play defense tactics

#### Long-Evans

There was a significant main effect of Playmate on the proportion of attacks defended and evaded, and role reversals (Table 8), with more attacks being defended, evaded, and resulting in more role reversals by familiar versus novel playmates (Fig. 4A, 4C, 4G).

**Fig. 4.**
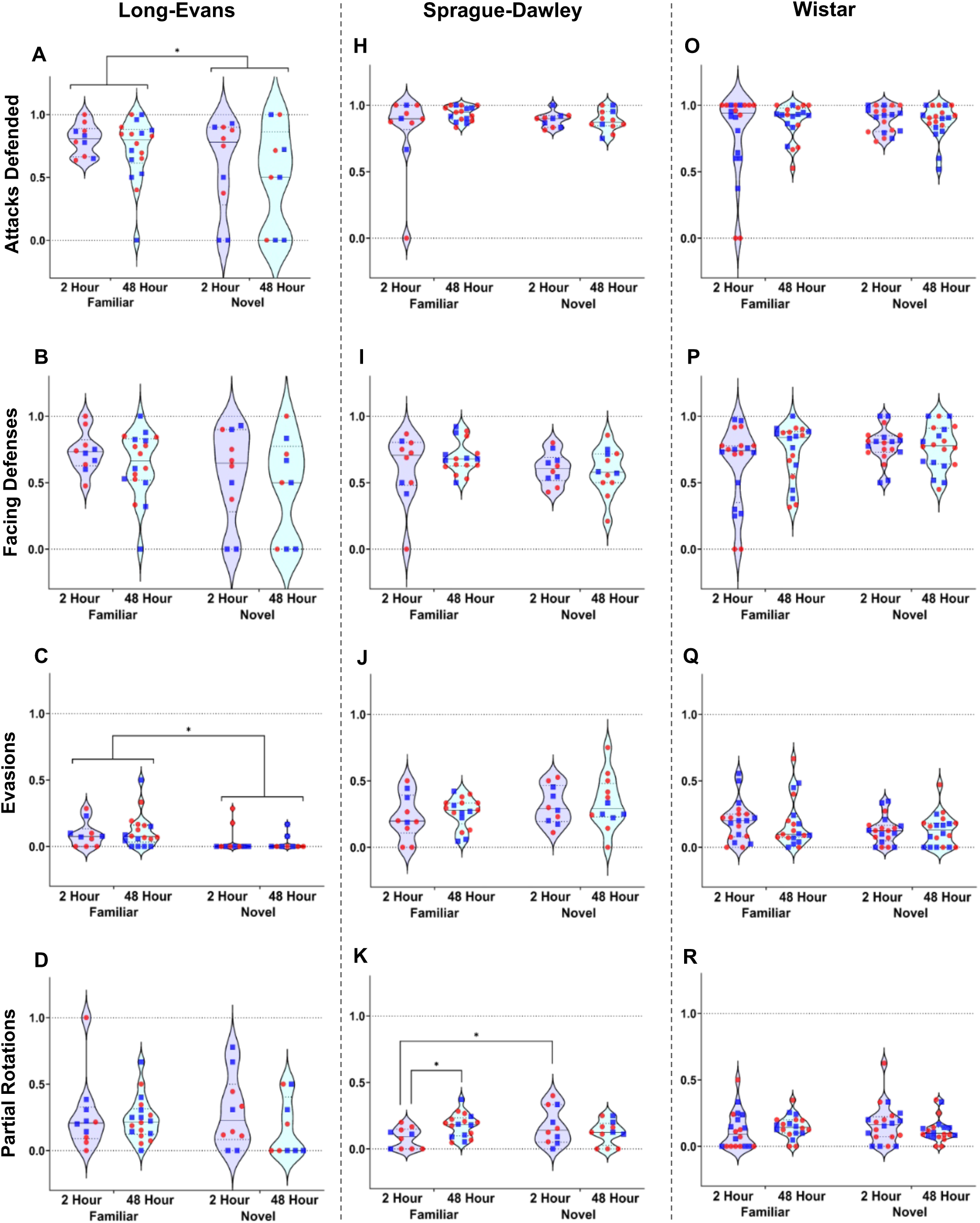
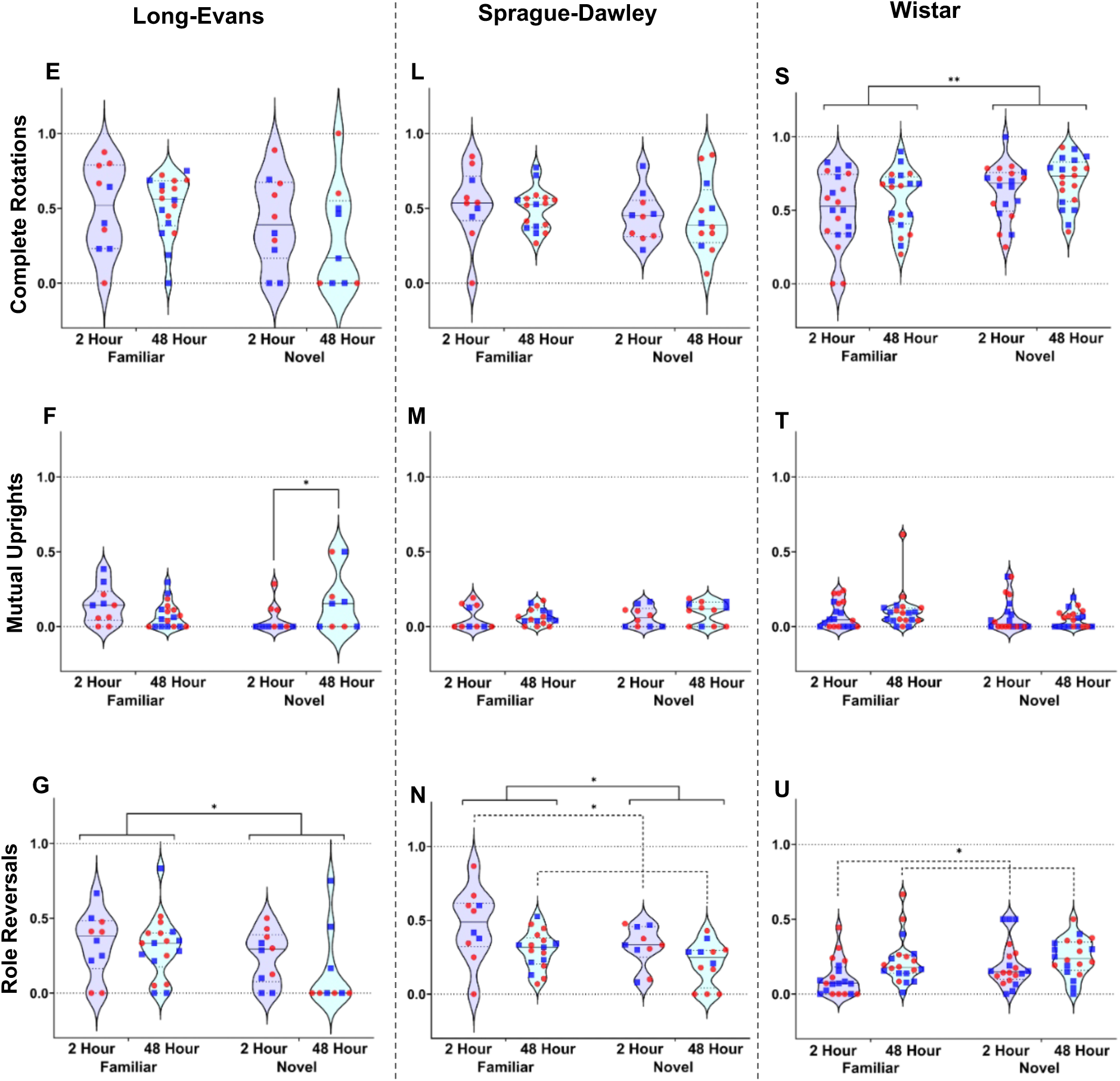
Experimental condition comparisons per rat strain for social play defense tactics of the playmates in response to nape attacks by the experimental rat – Most notably are more role reversals by familiar playmates in Long-Evans and Sprague-Dawley and fewer role reversals by Sprague-Dawley but more role reversals by Wistar after 48h isolation. **A-G:** Long-Evans familiar playmates showed higher proportions of attacks defended **(A)**, evasions **(C)**, and role reversals **(G)** than novel playmates irrespective of time isolated by the experimental Long-Evans rats. Long-Evans novel playmates showed a higher proportion of mutual uprights **(F)**when experimental rats were isolated for 48h versus 2h. Long-Evans familiar and novel playmates showed a higher proportion of role reversals **(G)** when experimental rats were isolated for 2h versus 48h. There were no experimental condition differences on facing defenses (**B**), partial rotations (**D**) or complete rotations (**E**). **H-N:** There were no experimental condition differences for Sprague-Dawley playmates on attacks defended **(H),** facing defenses (**I**), evasions **(J),** complete rotations (**L**), or mutual uprights **(M)**. However, Sprague-Dawley familiar playmates exposed to an experimental Sprague-Dawley rat isolated for 2h showed a lower proportion of partial rotations **(K)** compared to novel playmates and compared to familiar playmates exposed to an experimental Sprague-Dawley rat isolated for 48h. Sprague-Dawley familiar playmates showed a higher proportion of role reversals **(N)** than novel playmates irrespective of time isolated by the experimental rats. Moreover, Sprague-Dawley playmates exposed to an experimental rat isolated for 2h showed a higher proportion of role reversals (**N**) than those exposed to an experimental rat isolated for 48h. **O-U:** There were no experimental condition differences for Wistar playmates on attacks defended **(O),** facing defenses (**P**), evasions **(Q),** partial rotations **(R),** or mutual uprights **(T)**. However, Wistar novel playmates showed a higher proportion of complete rotations **(S)** than familiar playmates, irrespective of isolation time. Wistar playmates exposed to an experimental Wistar rat isolated for 48h showed a higher proportion of role reversals **(U)** than playmates exposed to an experimental Wistar rat isolated for 2h. Behaviors are expressed as proportions. Males are represented by blue squares, females are represented by red circles; Three-way ANOVA analysis per strain to assess effects of Stimulus, Isolation, and Sex, followed by Bonferroni post hoc tests to assess interaction effects; * *p* < 0.05, ** *p* ≤ 0.01, *** *p* ≤ 0.001.

**Table 8.** Experimental Condition comparisons for social play defense tactics in playmate Long-Evans rats. Three-way ANOVA statistics and partial eta squared (η2) effect sizes for the effects of Playmate (Familiar, Novel), Isolation (2h, 48h), and Sex on the proportion of social play tactic behaviors of playmate male and female juvenile Long-Evans rats in the social play test. Significant effects and their corresponding behaviors are indicated in **bold**.

|  | Playmate | Isolation | Playmate x Isolation | Sex |
| --- | --- | --- | --- | --- |
| **Attacks defended** | **$F_{(1,39)} = 5.27$,**<br>**$p = 0.027$**<br>**$\eta_p^2 = 0.12$** | $F_{(1,39)} = 0.99$,<br>$p = 0.326$ | $F_{(1,39)} = 0.02$,<br>$p = 0.878$ | $F_{(1,39)} = 2.00$,<br>$p = 0.166$ |
| Facing defenses | $F_{(1,39)} = 3.20$,<br>$p = 0.081$ | $F_{(1,39)} = 0.96$,<br>$p = 0.334$ | $F_{(1,39)} = 0.00$,<br>$p = 0.996$ | $F_{(1,39)} = 1.86$,<br>$p = 0.181$ |
| **Evasions** | **$F_{(1,39)} = 4.77$,**<br>**$p = 0.035$**<br>**$\eta_p^2 = 0.11$** | $F_{(1,39)} = 0.00$,<br>$p = 0.965$ | $F_{(1,39)} = 0.40$,<br>$p = 0.533$ | $F_{(1,39)} = 0.11$,<br>$p = 0.742$ |
| **Mutual uprights** | $F_{(1,39)} = 0.06$,<br>$p = 0.806$ | $F_{(1,39)} = 0.19$,<br>$p = 0.663$ | **$F_{(1,39)} = 7.23$,**<br>**$p = 0.011$**<br>**$\eta_p^2 = 0.16$** | $F_{(1,39)} = 0.18$,<br>$p = 0.676$ |
| **Complete rotations** | $F_{(1,39)} = 2.69$,<br>$p = 0.109$ | $F_{(1,39)} = 0.15$,<br>$p = 0.701$ | $F_{(1,39)} = 0.69$,<br>$p = 0.412$ | **$F_{(1,39)} = 6.50$,**<br>**$p = 0.015$**<br>**$\eta_p^2 = 0.14$** |
| Partial rotations | $F_{(1,39)} = 0.08$,<br>$p = 0.776$ | $F_{(1,39)} = 1.05$,<br>$p = 0.311$ | $F_{(1,39)} = 0.45$,<br>$p = 0.508$ | $F_{(1,39)} = 0.28$,<br>$p = 0.603$ |
| **Role reversals** | **$F_{(1,39)} = 4.73$,**<br>**$p = 0.036$**<br>**$\eta_p^2 = 0.11$** | $F_{(1,39)} = 1.44$,<br>$p = 0.237$ | $F_{(1,39)} = 0.29$,<br>$p = 0.593$ | $F_{(1,39)} = 0.95$,<br>$p = 0.336$ |

There was a significant Playmate x Isolation effect on the proportion of mutual uprights (Table 8). Bonferroni post hoc test showed that Long-Evans playmates in the 48h-Novel condition engaged in more mutual uprights compared to those in the 2h-Novel condition (Table 9, Fig. 4F).

**Table 9.** Posthoc tests for experimental condition comparisons for social play defense tactics in playmate Long-Evans rats. Bonferroni post hoc statistics and partial eta squared (η2) effect sizes for the interaction effects of Playmate (Familiar, Novel) and Isolation (2h, 48h) on the proportion of social play tactic behaviors of playmate male and female juvenile Long-Evans rats in the social play test. Significant effects and their corresponding behaviors are indicated in **bold**.

|  | 48-Hour Familiar<br>vs |  | 2-Hour Novel<br>vs |  |
| --- | --- | --- | --- | --- |
|  | 2-Hour Familiar | 48-Hour Novel | 2-Hour Familiar | 48-Hour Novel |
| **Mutual**<br>**uprights** | $F_{(1,43)} = 1.87,$<br>$p = 0.179$ | $F_{(1,43)} = 3.13,$<br>$p = 0.084$ | $F_{(1,43)} = 2.76,$<br>$p = 0.104$ | **$F_{(1,43)} = 4.07,$**<br>**$p = 0.050$**<br>**$\eta_p^2 = 0.09$** |

There was a significant main effect of Sex on the proportion of complete rotations (Table 8), with more complete rotations used by females than males. There was a significant effect of Playmate x Isolation x Sex on role reversals (Suppl. Table 8). Bonferroni tests showed that females engaged in more role reversals in the 48h-Familiar condition compared to the 48h-Novel condition (Suppl. Table 9). Moreover, females engaged in more role reversals in the 2h-Novel condition than 48h-Novel condition. Males engaged in more role reversals in the 2h-Familiar condition compared to the 2h-Novel condition.

#### Sprague-Dawley

There were significant main effects of Playmate and Isolation on the proportion of role reversals, with more role reversals occurring between familiar versus novel playmates and after 2h versus 48h isolation (Fig. 4N; Table 10).

**Table 10.** Experimental Condition comparisons for social play defense tactics in playmate Sprague-Dawley rats. Three-way ANOVA statistics and partial eta squared (η2) effect sizes for the effects of Playmate (Familiar, Novel), Isolation (2h, 48h), and Sex on the proportion of social play tactic behaviors of playmate male and female juvenile Sprague-Dawley rats in the social play test. Significant effects and their corresponding behaviors are indicated in **bold**.

|  | Playmate | Isolation | Playmate x Isolation | Sex |
| --- | --- | --- | --- | --- |
| Attacks defended | $F_{(1,41)} = 0.03$,<br>$p = 0.873$ | $F_{(1,41)} = 1.30$,<br>$p = 0.260$ | $F_{(1,41)} = 1.43$,<br>$p = 0.239$ | $F_{(1,41)} = 0.19$,<br>$p = 0.662$ |
| Facing defenses | $F_{(1,41)} = 0.64$,<br>$p = 0.430$ | $F_{(1,41)} = 0.81$,<br>$p = 0.374$ | $F_{(1,41)} = 0.90$,<br>$p = 0.350$ | $F_{(1,41)} = 0.54$,<br>$p = 0.468$ |
| Evasions | $F_{(1,41)} = 1.50$,<br>$p = 0.227$ | $F_{(1,41)} = 0.00$,<br>$p = 0.986$ | $F_{(1,41)} = 0.01$,<br>$p = 0.945$ | $F_{(1,41)} = 0.08$,<br>$p = 0.784$ |
| Mutual uprights | $F_{(1,41)} = 0.86$,<br>$p = 0.358$ | $F_{(1,41)} = 1.16$,<br>$p = 0.288$ | $F_{(1,41)} = 0.22$,<br>$p = 0.638$ | $F_{(1,41)} = 0.00$,<br>$p = 0.962$ |
| Complete rotations | $F_{(1,41)} = 1.24$,<br>$p = 0.273$ | $F_{(1,41)} = 0.07$,<br>$p = 0.797$ | $F_{(1,41)} = 0.07$,<br>$p = 0.788$ | $F_{(1,41)} = 0.71$,<br>$p = 0.403$ |
| **Partial rotations** | $F_{(1,41)} = 0.72$,<br>$p = 0.400$ | $F_{(1,41)} = 0.92$,<br>$p = 0.343$ | **$F_{(1,41)} = 4.63$,**<br>**$p = 0.037$**<br>**$\eta_p^2 = 0.10$** | $F_{(1,41)} = 0.02$,<br>$p = 0.888$ |
| **Role reversals** | **$F_{(1,41)} = 4.23$,**<br>**$p = 0.046$**<br>**$\eta_p^2 = 0.09$** | **$F_{(1,41)} = 6.47$,**<br>**$p = 0.015$**<br>**$\eta_p^2 = 0.14$** | $F_{(1,41)} = 0.54$,<br>$p = 0.465$ | $F_{(1,41)} = 0.37$,<br>$p = 0.548$ |

There was a significant Playmate x Isolation effect on the proportion of partial rotation (Fig. 4K; Table 10). Bonferroni post hoc tests showed that Sprague-Dawley playmates in the 2h-Familiar condition engaged in fewer partial rotations compared to the 48h-Familiar and 2h-Novel conditions (Table 11).

**Table 11.** Posthoc tests for experimental condition comparisons for social play defense tactics in playmate Sprague-Dawley rats. Bonferroni post hoc statistics and partial eta squared (η2) effect sizes for the interaction effects of Playmate (Familiar, Novel) and Isolation (2h, 48h) on the proportion of social play tactic behaviors of playmate male and female juvenile Sprague-Dawley rats in the social play test. Significant effects and their corresponding behaviors are indicated in **bold**.

|  | 48-Hour Familiar<br>vs |  | 2-Hour Novel<br>vs |  |
| --- | --- | --- | --- | --- |
|  | 2-Hour Familiar | 48-Hour Novel | 2-Hour Familiar | 48-Hour Novel |
| **Partial rotations** | **$F_{(1,45)} = 5.74,$**<br>**$p = 0.021$**<br>**$\eta_p^2 = 0.11$** | $F_{(1,45)} = 1.92,$<br>$p = 0.173$ | **$F_{(1,45)} = 4.22,$**<br>**$p = 0.046$**<br>**$\eta_p^2 = 0.09$** | $F_{(1,45)} = 1.29,$<br>$p = 0.262$ |

There were no significant main effects of Sex or interaction effects with Sex on any play defense behavior analyzed (Table 10).

#### Wistar

There was a significant main effect of Playmate on the proportion of complete rotations, with more complete rotations occurring by novel versus familiar playmates (Fig. 4S; Table 12). There was a significant main effect of Isolation on the proportion of role reversals, with more role reversals occurring in the 48h isolation versus 2h isolation condition (Fig. 4U; Table 12). There were no other significant main or interaction effects.

**Table 12.** Experimental Condition comparisons for social play defense tactics in playmate Wistar rats. Three-way ANOVA statistics and partial eta squared (η2) effect sizes for the effects of Playmate (Familiar, Novel), Isolation (2h, 48h), Sex on the proportion of social play tactic behaviors of playmate male and female juvenile Wistar rats in the social play test. Significant effects are indicated in **bold**.

|  | Playmate | Isolation | Playmate x Isolation | Sex |
| --- | --- | --- | --- | --- |
| Attacks defended | $F_{(1,72)} = 1.56,$<br>$p = 0.216$ | $F_{(1,72)} = 0.77,$<br>$p = 0.384$ | $F_{(1,72)} = 1.62,$<br>$p = 0.208$ | $F_{(1,72)} = 0.12,$<br>$p = 0.733$ |
| Facing defenses | $F_{(1,72)} = 3.81,$<br>$p = 0.055$ | $F_{(1,72)} = 0.71,$<br>$p = 0.404$ | $F_{(1,72)} = 1.36,$<br>$p = 0.248$ | $F_{(1,72)} = 0.00,$<br>$p = 0.955$ |
| Evasions | $F_{(1,72)} = 3.40,$<br>$p = 0.069$ | $F_{(1,72)} = 0.05,$<br>$p = 0.823$ | $F_{(1,72)} = 0.07,$<br>$p = 0.791$ | $F_{(1,72)} = 0.02,$<br>$p = 0.879$ |
| Mutual uprights | $F_{(1,72)} = 1.26,$<br>$p = 0.266$ | $F_{(1,72)} = 0.00,$<br>$p = 0.952$ | $F_{(1,72)} = 1.43,$<br>$p = 0.236$ | $F_{(1,72)} = 2.70,$<br>$p = 0.105$ |
| **Complete rotations** | **$F_{(1,72)} = 7.21,$**<br>**$p = 0.009$**<br>**$\eta_p^2 = 0.09$** | $F_{(1,72)} = 1.92,$<br>$p = 0.170$ | $F_{(1,72)} = 0.00,$<br>$p = 0.961$ | $F_{(1,72)} = 2.31,$<br>$p = 0.133$ |
| Partial rotations | $F_{(1,72)} = 0.20,$<br>$p = 0.654$ | $F_{(1,72)} = 0.08,$<br>$p = 0.786$ | $F_{(1,72)} = 1.58,$<br>$p = 0.214$ | $F_{(1,72)} = 0.24,$<br>$p = 0.626$ |
| **Role reversals** | $F_{(1,72)} = 3.56,$<br>$p = 0.063$ | **$F_{(1,72)} = 6.67,$**<br>**$p = 0.012$**<br>**$\eta_p^2 = 0.08$** | $F_{(1,72)} = 1.09,$<br>$p = 0.300$ | $F_{(1,72)} = 2.54,$<br>$p = 0.115$ |

### 3. Strain comparisons per condition for social play behavior

#### 2h-Familiar Condition

There were no significant strain differences on the duration of social play, number of nape attacks, and number of pins. There was a significant strain difference on the number of supine poses (Table 13) with Bonferroni post hoc tests revealing that Long-Evans showed more supine poses than Wistar (*p* = 0.006; Fig. 5D).

**Fig. 5.**
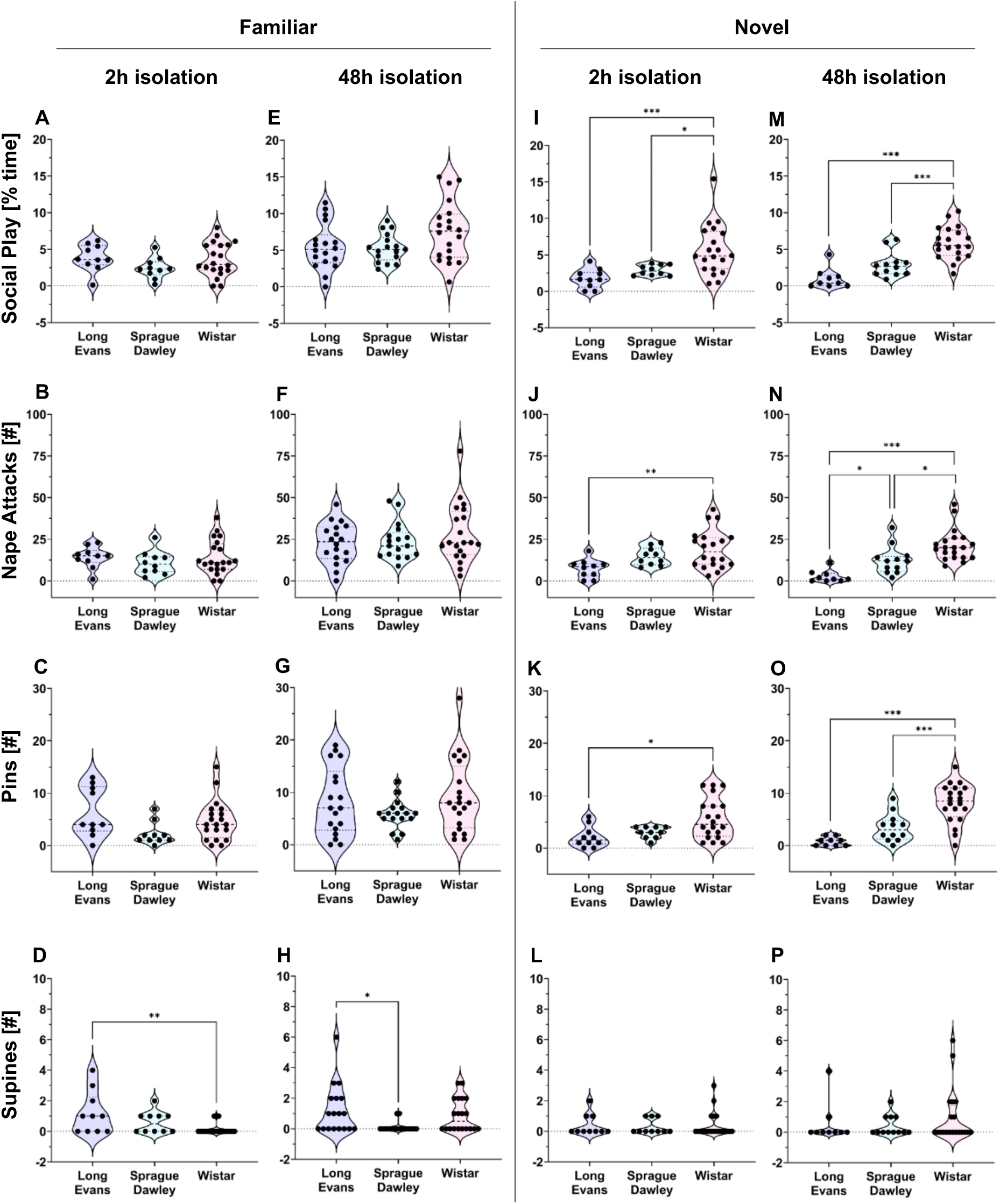
Rat strain comparisons for social play behaviors per experimental condition - Strain differences are most prevalent when rats were exposed to a novel playmate. **A-D:** There were no strain differences on social play behaviors when exposed to a familiar playmate after 2h isolation, except for fewer supines in Wistar compared to Long Evans **(D)**. **E-H:** There were no strain differences on social play behaviors when exposed to a familiar playmate after 48h isolation, except for fewer supines in Sprague-Dawley compared to Long Evans **(H)**. **I-K:** Long Evans played less than Wistar when exposed to a novel playmate after 2h isolation, as shown by shorter duration of social play (**I)**, fewer nape attacks (**J**), and fewer pins (**K**). **I:** Sprague Dawley played less than Wistar when exposed to a novel playmate after 2h isolation. **M-O:** Long Evans and Sprague-Dawley played less than Wistar when exposed to a novel playmate after 48h isolation, as shown by shorter duration of social play (**M)**, fewer nape attacks (**N**), and fewer pins (**O**). **N:** Long-Evans had fewer nape attacks than Sprague Dawley when exposed to a novel playmate after 48h isolation. **L, P:** There were no strain differences on supines when exposed to a novel playmate after 2h or 48h isolation. One-way ANOVA analysis per condition followed by Bonferroni post hoc tests to assess strain differences; * *p* < 0.05, ** *p* ≤ 0.01, *** *p* ≤ 0.001.

**Table 13.** Strain comparisons for social play behaviors of experimental rats in each of the four experimental conditions. One-way ANOVA statistics and partial eta squared (η2) effect sizes for the effect of Strain on behavior of experimental male and female juvenile Long-Evans, Sprague-Dawley, and Wistar rats in the social play test. Significant effects are indicated in **bold**.

|  | 2-Hour Familiar | 48-Hour Familiar | 2-Hour Novel | 48-Hour Novel |
| --- | --- | --- | --- | --- |
| **Social play**<br>**(% Time)** | $F_{(2,37)} = 1.60,$<br>$p = 0.215$ | $F_{(2,52)} = 3.06,$<br>$p = 0.055$<br>$\eta_p^2 = 0.11$ | **$F_{(2,37)} = 8.87,$**<br>**$p < 0.001$**<br>**$\eta_p^2 = 0.32$** | **$F_{(2,38)} = 23.0,$**<br>**$p < 0.001$**<br>**$\eta_p^2 = 0.55$** |
| **Nape attacks (#)** | $F_{(2,37)} = 0.59,$<br>$p = 0.560$ | $F_{(2,52)} = 1.01,$<br>$p = 0.372$ | **$F_{(2,37)} = 5.53,$**<br>**$p = 0.008$**<br>**$\eta_p^2 = 0.23$** | **$F_{(2,38)} = 16.6,$**<br>**$p < 0.001$**<br>**$\eta_p^2 = 0.47$** |
| **Pins (#)** | $F_{(2,37)} = 3.08,$<br>$p = 0.058$ | $F_{(2,52)} = 1.61,$<br>$p = 0.209$ | **$F_{(2,37)} = 5.66,$**<br>**$p = 0.007$**<br>**$\eta_p^2 = 0.23$** | **$F_{(2,38)} = 21.0,$**<br>**$p < 0.001$**<br>**$\eta_p^2 = 0.52$** |
| **Supine poses (#)** | **$F_{(2,37)} = 5.60,$**<br>**$p = 0.008$**<br>**$\eta_p^2 = 0.23$** | **$F_{(2,52)} = 4.39,$**<br>**$p = 0.017$**<br>**$\eta_p^2 = 0.14$** | $F_{(2,37)} = 0.07,$<br>$p = 0.931$ | $F_{(2,38)} = 0.60,$<br>$p = 0.554$ |

#### 48h-Familiar Condition

There were no significant strain differences on the duration of social play, number of nape attacks, and number of pins. There was a significant strain difference on the number of supine poses (Table 13), with Bonferroni post hoc tests revealing that Long-Evans showed more supine poses than Sprague-Dawley (*p* = 0.017; Fig. 5H).

#### 2h-Novel Condition

There were significant strain differences on the duration of social play, number of nape attacks, and number of pins (Table 13), with Bonferroni post hoc tests revealing that Wistar rats showed more social play than Long-Evans (*p* = 0.001) and Sprague-Dawley (*p* = 0.026; Fig. 5I) and more nape attacks (*p* = 0.006) and pins (*p* = 0.012) than Long-Evans (Fig. 5J, 5K). There were no significant strain differences on the number of supine poses.

#### 48h-Novel Condition

There were significant strain differences on the duration of social play, number of nape attacks, and number of pins (Table 13). Bonferroni post hoc tests revealed that Wistar showed more social play (Fig. 5M), more nape attacks (Fig. 5N) and more pins (Fig. 5O) than Long-Evans (*p* < 0.001 for social play, nape attacks, and pins) and Sprague-Dawley (*p* < 0.001 for social play and pins; *p* = 0.016 for nape attacks). In addition, Sprague-Dawley showed more nape attacks than Long-Evans (*p* = 0.032; Fig. 5N). There were no strain differences on the number of supine poses.

### 4. Strain comparisons per condition for social play defense tactics

#### 2h-Familiar Condition

There were significant strain differences on the proportion of role reversals (Table 14), with Bonferroni post hoc tests revealing that both Long-Evans (*p* = 0.010; Fig. 6G) and Sprague-Dawley (*p* < 0.001) playmates engaged in more role reversals than Wistar playmates. No other strain differences in defensive tactics were found.

**Fig. 6.**
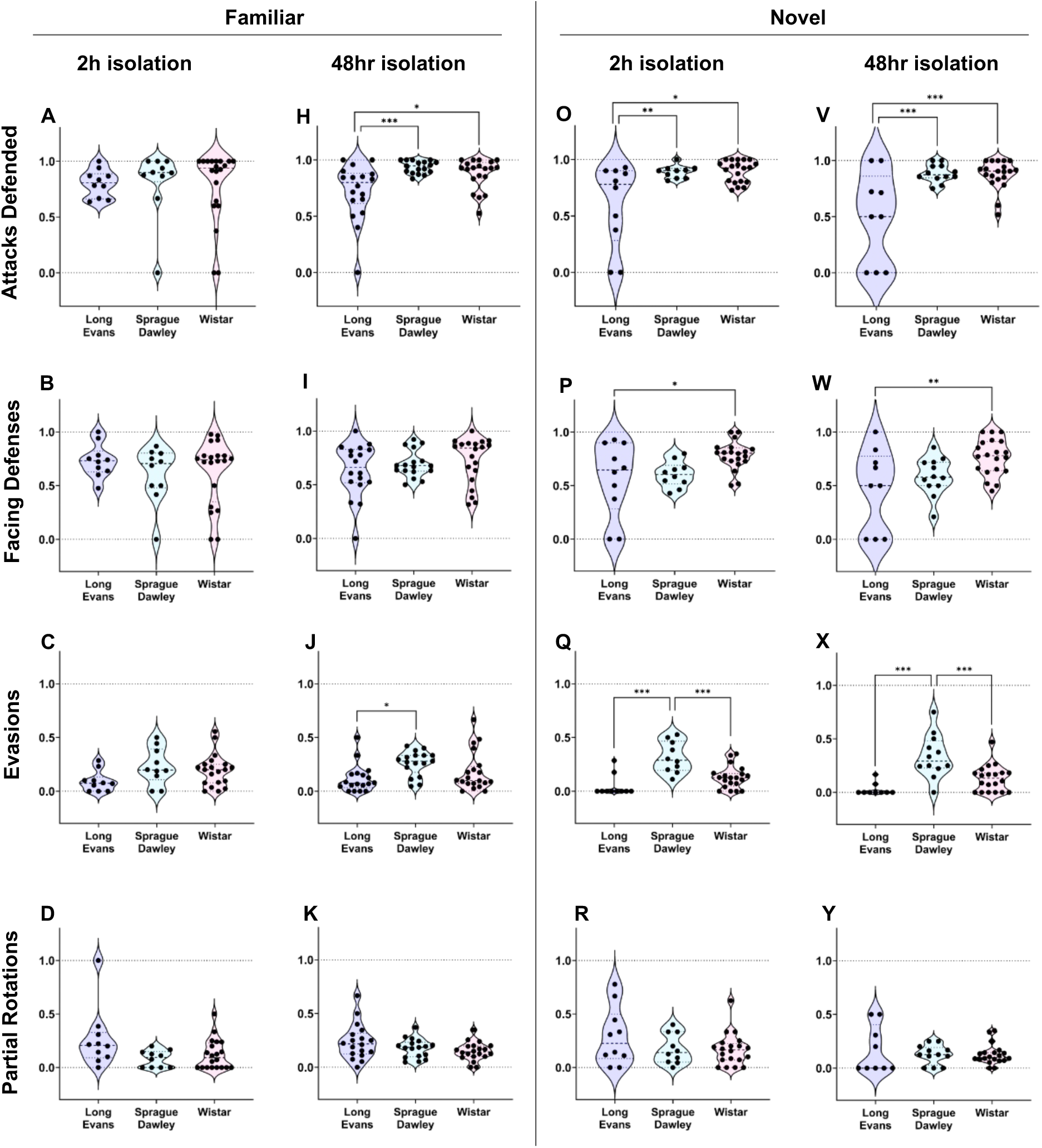
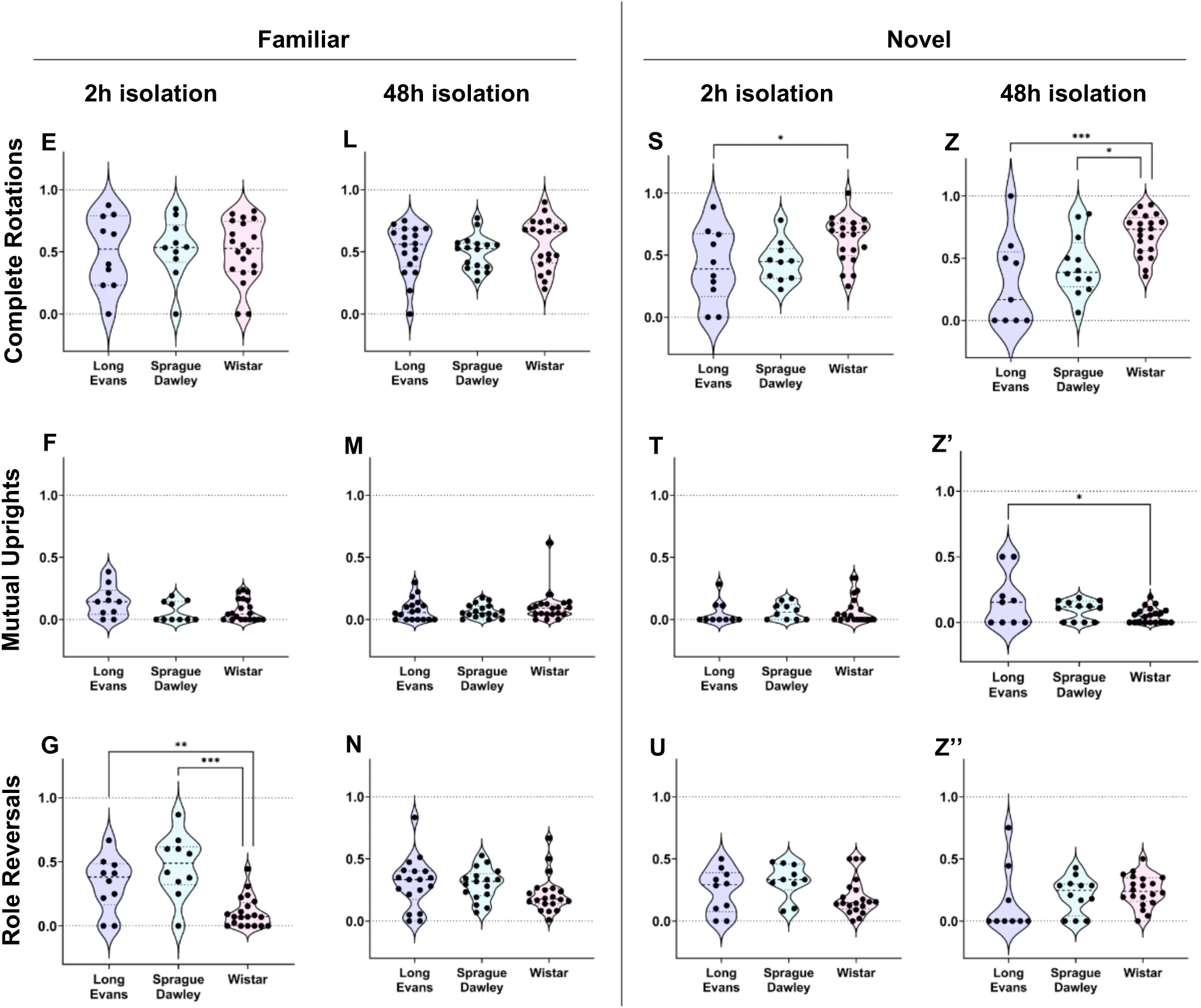
Rat strain comparisons per experimental condition for social play defense tactics of the playmates in response to nape attacks by the experimental rat – Most notable are strain differences with novel playmates with more attacks defended by Sprague-Dawley and Wistar, more evasions by Sprague-Dawley and more facing defenses and complete rotations by Wistar. **A-G:** When experimental rats were exposed to a familiar playmate after 2h isolation, there were no strain differences on the proportions of attacks defended (**A**), facing defenses (**B**), evasions (**C**), partial rotations (**D**), complete rotations (**E**), or mutual uprights (**F**) by the playmates. However, Wistar playmates showed fewer role reversals (**G**) than Long-Evans and Sprague-Dawley playmates. **H-N:** When experimental rats were exposed to a familiar playmate after 48h isolation, Long Evans playmates showed fewer defended attacks (**H**) than Sprague-Dawley and Wistar playmates and fewer evasions (**J**) than Sprague-Dawley playmates. No strain differences were found for the other defense tactics. **O-U:** When experimental rats were exposed to a novel playmate after 2h isolation, Long Evans playmates showed fewer defended attacks (**O**) than Sprague-Dawley and Wistar playmates and fewer facing defenses (**P**) and complete rotations (**S**) than Wistar playmates. Sprague-Dawley playmates showed more evasions (**Q**) than Long-Evans and Wistar playmates. No strain differences were found for the other defense tactics. **V-Z’’:** When experimental rats were exposed to a novel playmate after 48h isolation, Long Evans playmates showed fewer defended attacks (**V**) but more mutual uprights (**Z’**) than Sprague-Dawley and Wistar playmates. Wistar playmates showed more complete rotations (**Z**) than Long-Evans and Sprague-Dawley playmates. No strain differences were found for the other defense tactics. Behaviors are expressed as proportions. One-way ANOVA analyses per experimental condition followed by Bonferroni post hoc tests to assess strain differences; * *p* < 0.05, ** *p* ≤ 0.01, *** *p* ≤ 0.001.

**Table 14.** Strain comparisons for social play defense tactics of playmate rats in each of the four experimental conditions. One-way ANOVA statistics and partial eta squared (η2) effect sizes for the effect of Strain on the proportion of social play tactic behaviors of playmate male and female juvenile Long-Evans, Sprague-Dawley, and Wistar rats in the social play test. Significant effects are indicated in **bold**.

|  | 2-Hour Familiar | 48-Hour Familiar | 2-Hour Novel | 48-Hour Novel |
| --- | --- | --- | --- | --- |
| **Attacks defended** | $F_{(2,37)} = 0.04$,<br>$p = 0.961$ | **$F_{(2,37)} = 7.84$,**<br>**$p = 0.001$**<br>**$\eta_p^2 = 0.23$** | **$F_{(2,37)} = 8.20$,**<br>**$p = 0.001$**<br>**$\eta_p^2 = 0.31$** | **$F_{(2,37)} = 11.6$,**<br>**$p < 0.001$**<br>**$\eta_p^2 = 0.38$** |
| **Facing defenses** | $F_{(2,37)} = 0.68$,<br>$p = 0.513$ | $F_{(2,37)} = 0.95$,<br>$p = 0.394$ | **$F_{(2,37)} = 4.76$,**<br>**$p = 0.014$**<br>**$\eta_p^2 = 0.20$** | **$F_{(2,37)} = 5.74$,**<br>**$p = 0.007$**<br>**$\eta_p^2 = 0.23$** |
| **Evasions** | $F_{(2,37)} = 2.58$,<br>$p = 0.089$ | **$F_{(2,37)} = 4.37$,**<br>**$p = 0.018$**<br>**$\eta_p^2 = 0.14$** | **$F_{(2,37)} = 15.2$,**<br>**$p < 0.001$**<br>**$\eta_p^2 = 0.45$** | **$F_{(2,37)} = 14.1$,**<br>**$p < 0.001$**<br>**$\eta_p^2 = 0.43$** |
| **Mutual uprights** | $F_{(2,37)} = 2.29$,<br>$p = 0.115$ | $F_{(2,37)} = 0.71$,<br>$p = 0.495$ | $F_{(2,37)} = 0.24$,<br>$p = 0.786$ | **$F_{(2,37)} = 3.50$,**<br>**$p = 0.040$**<br>**$\eta_p^2 = 0.16$** |
| **Complete rotations** | $F_{(2,37)} = 0.03$,<br>$p = 0.973$ | $F_{(2,37)} = 0.84$,<br>$p = 0.438$ | **$F_{(2,37)} = 4.36$,**<br>**$p = 0.020$**<br>**$\eta_p^2 = 0.19$** | **$F_{(2,37)} = 9.41$,**<br>**$p < 0.001$**<br>**$\eta_p^2 = 0.33$** |
| Partial rotations | $F_{(2,37)} = 3.12$,<br>$p = 0.056$ | $F_{(2,37)} = 2.85$,<br>$p = 0.067$ | $F_{(2,37)} = 1.64$,<br>$p = 0.208$ | $F_{(2,37)} = 0.38$,<br>$p = 0.687$ |
| **Role reversals** | **$F_{(2,37)} = 14.3$,**<br>**$p < 0.001$**<br>**$\eta_p^2 = 0.44$** | $F_{(2,37)} = 1.62$,<br>$p = 0.208$ | $F_{(2,37)} = 2.17$,<br>$p = 0.129$ | $F_{(2,37)} = 0.92$,<br>$p = 0.409$ |

#### 48h-Familiar Condition

There were significant strain differences on the proportion of attacks defended and evaded (Table 14). Bonferroni post hoc tests revealed that both Sprague-Dawley (*p* = 0.001; Fig. 6H) and Wistar (*p* = 0.017) playmates defended more playful attacks than Long-Evans playmates. Additionally, Sprague-Dawley playmates were more likely to evade a playful attack than Long-Evans playmates (*p* = 0.014; Fig. 6J). There were no other strain differences in defensive tactics.

#### 2h-Novel Condition

There were significant strain differences on the proportion of attacks defended, facing defenses, evasions, and complete rotations (Table 14). Bonferroni post hoc tests revealed that both Sprague-Dawley (*p* = 0.007; Fig. 6O) and Wistar (*p* = 0.001) playmates defended more playful attacks than Long-Evans playmates. Wistar playmates also engaged in more facing defenses compared to Long-Evans playmates (*p* = 0.027; Fig. 6P). Sprague-Dawley playmates evaded playful attacks more than both Long-Evans (*p* < 0.001; Fig. 6Q) and Wistar (*p* < 0.001) playmates. Finally, Wistar playmates defended playful attacks with complete rotations significantly more than Long-Evans playmates (*p* = 0.041; Fig. 6S).

#### 48h-Novel Condition

There were significant strain differences on the proportion of attacks defended, facing defenses, evasions, mutual uprights, and complete rotations (Table 14). Bonferroni post hoc tests revealed that both Sprague-Dawley (*p* < 0.001; Fig. 6V) and Wistar (*p* < 0.001) playmates defended more attacks than Long-Evans playmates. Wistar playmates also engaged in more facing defenses than Long-Evans playmates (*p* = 0.008; Fig. 6W). Sprague-Dawley playmates evaded playful attacks more than Long-Evans (*p* < 0.001; Fig. 6X) and Wistar (*p* < 0.001) playmates. Additionally, Wistar playmates engaged in more complete rotations than both Long-Evans (*p* < 0.001; Fig. 6Z) and Sprague-Dawley (*p* = 0.021) playmates. Finally, Long-Evans playmates engaged in mutual uprights more than Wistar playmates (*p* = 0.036; Fig. 6Z’).

## DISCUSSION

We found that playmate familiarity (familiar versus novel) and social isolation length (2h versus 48h) influenced social play levels and tactics in juvenile rats, but did so differently for each of the three rat strains. Long-Evans rats played significantly more with a familiar than with a novel playmate, irrespective of time isolated, demonstrating that playmate familiarity influences social play for this strain. Sprague-Dawley rats played more in the 48h-Familiar condition compared to both the 2h-Familiar condition and 48h-Novel condition, demonstrating that for this strain a combination of longer social isolation with exposure to a familiar playmate facilitates social play expression. Wistar rats played less in the 2h-Familiar condition compared to both the 48h- Familiar condition and the 2h-Novel condition, demonstrating that either a longer isolation from a familiar playmate or exposure to a novel playmate facilitates social play in this strain. Moreover, strain differences in social play levels emerged when rats were exposed to a novel playmate. Here, Wistar rats played more with a novel playmate than Long-Evans and Sprague-Dawley rats, irrespective of isolation time. Playmate familiarity influenced social play tactics in Long-Evans rats with more attacks defended, more evasions, and more role reversals by a familiar than a novel playmate. A combination of playmate familiarity and social isolation length influences social play tactics in Sprague-Dawley with more role reversals by a familiar versus a novel playmate and after 2h versus 48h isolation. Isolation length influenced social play tactics in Wistar with more role reversals after 48h isolation. Additionally, strain differences in social play tactics were most prevalent in novel playmates with more attacks defended by Sprague-Dawley and Wistar versus Long-Evans. This was followed by more evasions by Sprague-Dawley playmates. In contrast, Wistar playmates turned to face their playmate attacker more and showed more complete rotations, thereby lengthening body contact and wrestling bouts. Across the three rat strains, the effects of playmate familiarity and social isolation length on social play levels and tactics were similar across sex. Taken together, Long-Evans showed more and better quality (indicated by role reversals) of social play with familiar playmates. Sprague-Dawley also played more with familiar playmates and showed better quality of social play with familiar playmates after a short isolation. Wistar played equally well with familiar and novel playmates, but the quality of play was higher after a longer isolation time. Given that the majority of social play research is conducted in rats, our findings underscore the importance of taking into account variations in external conditions and type of strain when interpreting outcomes. Future research could determine the neuronal mechanisms mediating strain- and condition-induced variations in social play levels and tactics.

### 1. Effects of playmate familiarity and social isolation length on social play levels

#### Long-Evans play more with familiar than novel playmates, irrespective of isolation time

Long-Evans rats have been commonly used to study the ontogeny, function, and neural basis of social play behavior using a wide variety of social play paradigms (Ham and Pellis, 2024; Ku et al., 2016; Parent and Meaney, 2008; Stark et al., 2022; Takahashi and Lore, 1983). Even so, and to the best of our knowledge, our study is the first demonstrating that juvenile male and female Long-Evans rats played more with a familiar than with a novel playmate, irrespective of a shorter (2h) or longer (48h) isolation time. A previous study showed that juvenile male and female Long-Evans rats had similar play levels when exposed to an unfamiliar sibling (that they had not seen since weaning) versus a sibling cagemate (two-way ANOVA, main effect of play partner: F_(1, 36)_ = 0.87, p = 0.36, statistical analysis provided by Dr. Katherina Northcutt based on Northcutt and Nwankwo (2018). A notable difference between this study and our study is that the rats in the latter study were not socially isolated prior to social play testing. Social isolation typically increases social play levels, which is driven, in part, by increased social motivation (Achterberg et al., 2023; Ikemoto and Panksepp, 1992; Vanderschuren et al., 2016). Indeed, several studies have demonstrated that a longer duration of social isolation (varying from no social isolation up to 15 days) increased social play levels in juvenile male and female Long-Evans rats (Panksepp, 1981; Panksepp and Beatty, 1980; Thor and Holloway, 1984). This finding was independent of the play partner’s sex or familiarity. We also found that a longer isolation time (48h vs 2h) increased social play levels in juvenile male and female Long-Evans rats. However, this was true when rats were exposed to a familiar playmate but not when they were exposed to a novel playmate. The lower levels of social play with a novel playmate could be due to social neophobia. Yet, adult female Long-Evans rats worked harder for access to a non-cagemate than a cagemate peer (Hackenberg et al., 2021). Moreover, adult male and female Long-Evans rats spent a similar time with a same-sex partner compared to a same-sex stranger in a 3h partner preference test (Beery and Shambaugh, 2021). However, neither study was performed in the home cage. Based on our own breeding experience with Long-Evans, reproductive success increased when the female was housed in the male’s home cage rather than in the female’s home cage. Taken together, it would be interesting to determine whether Long-Evans prefer to play with a familiar over a novel playmate when given the choice and whether this preference changes when social play testing would occur in a neutral versus a home cage.

#### Sprague-Dawley play most with a familiar playmate after a 48h isolation period

We found that social play in male and female Sprague-Dawley rats is driven by a combination of playmate familiarity and isolation time, with higher social play levels after a 48h isolation period and exposure to a familiar playmate. This finding is in line with another study showing more play-soliciting interactions (females only) and more social investigation (both sexes) when juvenile Sprague-Dawley rats were reunited after 24h isolation with a familiar as opposed to an unfamiliar peer in a neutral cage (Livia Terranova et al., 1999). Moreover, juvenile female, but not male, Sprague-Dawley rats played more with a familiar than a novel playmate in a neutral cage without prior isolation (Argue and McCarthy, 2015). Yet, juvenile male and female Sprague-Dawley rats had similar social play levels when exposed to an unfamiliar versus familiar sibling without prior isolation (two-way ANOVA, main effect of play partner: F_(1, 38)_ = 0.006, p = 0.936, statistical analysis provided by Dr. Katherina Northcutt based on Northcutt and Nwankwo (2018). Finally, single-housed, but not pair-housed, adult female (males were not tested) Sprague-Dawley rats earned more social interactions when lever-pressing for a familiar than for a novel partner (Chow et al., 2022). Despite the different paradigms used, these studies together provide evidence suggesting that Sprague-Dawley rats prefer interacting with a familiar over a novel conspecific, an effect that may remain consistent across the lifespan and may require a period of social isolation.

#### Wistar play least with a familiar playmate after a 2h isolation period

Male and female Wistar rats had significantly lower levels of social play in the 2h-Familiar condition compared to the 48h-Familiar condition. This suggests that the motivation to play with a familiar playmate in Wistar rats increases with a longer period of social isolation. A previous study showed that the number of pins increased with longer social isolation periods (i.e., 0, 1, 2, 3.5, 4, 6, 24 or 48 h; leveling off after 24h) in 4-week-old male Wistar rats when tested with a familiar rat (Niesink and Van Ree, 1989). This shows support that a relatively longer social isolation facilitates the expression of play behaviors with a familiar playmate in Wistar rats. Additionally, Wistar rats played significantly more with a novel than a familiar playmate after 2h isolation and played equally well with a novel and familiar playmate after 48h isolation. This finding is supported by a previous study showing that without prior social isolation, juvenile male and female Wistar rats played more with an unfamiliar sibling (that they had not seen since weaning) than with a sibling cagemate (two-way ANOVA, main effect of play partner: F_(1, 40)_ = 4.65, p = 0.037, no significant sex or interaction effects; statistical analysis provided by Dr. Katherina Northcutt based on Northcutt and Nwankwo (2018). To the best of our knowledge, no other studies compared social play levels between familiar and novel playmate exposure in Wistar rats. However, several studies compared social play levels in juvenile Wistar rats exposed to either a novel or familiar environment (Bredewold et al., 2014; Trezza and Vanderschuren, 2008; Vanderschuren et al., 1995b). When juvenile male Wistar rats were tested for social play with a familiar playmate in a novel versus a familiar test cage after a 3.5h isolation, they showed fewer pins in the first 5 min, but more pins in the next 5 min, leading to similar number of pins over the full length of the test (Trezza and Vanderschuren, 2008; Vanderschuren et al., 1995b). Likewise, social play behaviors were similar when juvenile male and female Wistar rats were exposed to a novel playmate in a novel versus familiar test cage after a 3-5 day isolation (Bredewold et al., 2014). These findings may show that novelty perse (novel playmate or novel test cage) does not negatively influence the expression of social play behaviors in Wistar rats. In contrast, a 3h isolation period is sufficient to reduce social novelty preference in a 3-chamber choice test by primarily increasing time spent with a cage mate in juvenile male and female Wistar rats (Smith et al., 2018, 2015). Taken together, it would be interesting to determine whether juvenile Wistar rats have a preference to play with a familiar over a novel playmate when given a choice and whether this changes depending on the social isolation time.

#### No sex differences in the effects of playmate familiarity or social isolation length on social play levels

We found that the effects of playmate familiarity and isolation time on social play levels were similar across sex for all three rat strains. There were also no sex differences in social play duration, number of nape attacks, and number of pins in Long-Evans and Sprague-Dawley rats. However, Sprague-Dawley males showed more supines and Wistar males showed higher levels of social play and more nape attacks than females. The lack of sex differences in the major elements of social play behavior in Long-Evans and Sprague-Dawley is supported by several other studies (Kisko et al., 2018; Lee et al., 2024; Northcutt and Nguyen, 2014; Northcutt and Nwankwo, 2018). These studies use a dyadic design, similar to our social play test. Yet, other studies reported sex differences in social play in Long-Evans (Meaney et al., 1983, 1981; Meaney and McEwen, 1986; Meaney and Stewart, 1981; Takahashi and Lore, 1983) and Sprague-Dawley (Jessen et al., 2010; Kurian et al., 2008; Marquardt et al., 2023; Olesen et al., 2005; Olioff and Stewart, 1978). All of these studies found higher social play levels in males compared to females. Most of these studies used a focal observation design with multiple rats in a cage and often mixed-sexed. Thus, sex differences in social play are not uniformly found, which led to the suggestion that variations in testing paradigms may explain the presence or absence of sex differences (Panksepp, 1981; Panksepp and Beatty, 1980; Thor and Holloway, 1984; VanRyzin et al., 2020). The same seems true for Wistar rats, with some studies reporting sex differences in social play (Birke and Sadler, 1983; Götz et al., 1991; Lukas and Wöhr, 2015; Lundberg et al., 2017), and other studies, including those from our lab, reporting no sex differences in social play (Bredewold et al., 2018, 2015; Northcutt and Nwankwo, 2018; Paul et al., 2014; Veenema et al., 2013). The inconsistency of finding sex differences in social play even in one lab, namely ours, might be worrisome. However, we observed that there were no significant sex differences in social play levels when each of the four experimental conditions was tested separately, suggesting that a high number of rats (i.e. n=40/sex) is required to find a sex difference in social play in Wistar rats using a dyadic design.

#### Strain differences in social play levels upon exposure to a novel, but not a familiar, playmate

Long-Evans, Sprague-Dawley and Wistar rats showed similar levels of social play when they were exposed to a familiar playmate. However, there were robust strain differences in social play levels when they were exposed to a novel playmate. In detail, Wistar played more (longer social play duration, more pins) with a novel playmate than Long-Evans and Sprague-Dawley, irrespective of isolation time. Moreover, Sprague-Dawley played more (longer social play duration, more nape attacks, more pins) with a novel playmate than Long-Evans after a 48h isolation period. Our findings are partly supported by Northcutt and Nwankwo (2018) reporting that Wistar rats played more with an unfamiliar playmate compared to Sprague-Dawley. However, none of these two strains differed in their play levels compared to Long-Evans (Northcutt and Nwankwo, 2018). Northcutt and Nwankwo (2018) did not isolate the experimental rats prior to testing, which may explain the lack of a difference in social play levels between Wistar and Long-Evans and between Sprague-Dawley and Long-Evans. In support, Sprague-Dawley showed more nape attacks than Long-Evans (regardless of playmate familiarity) after a 24h isolation (Himmler et al., 2014a). Moreover, Sprague-Dawley had higher levels of social play and more nape attacks with a novel playmate than Long-Evans rats after only a 10 minute isolation (Ku et al., 2016). Indeed, when male Long-Evans are playing in groups, and so can choose with whom they wish to play, they play more with novel conspecifics, playing little with their familiar cage mates (Ham and Pellis, 2023). These findings together suggest that all strains are motivated to play with novel partners, but among these three strains, the motivation to play with a novel playmate is the highest in Wistar rats while social isolation reduces, rather than enhances, the motivation to play in Long-Evans rats.

The observed strain differences in social play levels with a novel playmate may be due to strain differences in general and/or social neophobia. Both adult albino/Wistar and hooded/Long-Evans rats initially avoided eating from a novel food container when there was a familiar food container present (Mitchell, 1976), which suggests the presence of neophobia in both strains. However, albino/Wistar required less time than hooded/Long-Evans to eat equally from both containers (Mitchell, 1976), demonstrating less neophobia in Wistar compared to Long-Evans. Yet, potential strain differences in social neophobia are less clear. Juvenile male and female Wistar rats spend more time investigating a novel over a familiar conspecific in a 10-min social novelty preference test (Smith et al., 2015), suggesting low levels of social neophobia. Although similar studies are lacking for juvenile Long-Evans, adult Long-Evans rats spent more time in contact with a novel rat than they did with a familiar rat in a 5-min social novelty preference test (Fujihara et al., 2021). Lastly, juvenile-adolescent male (postnatal days 27-43) and adult male and female Sprague-Dawley rats spent more time investigating a novel over a familiar conspecific in a 10-min social novelty preference test (Hughes et al., 2020b, 2020a; Win-Shwe et al., 2021). These findings do not support the suggestion that strain differences in social play levels with a novel playmate are due to strain differences in social neophobia.

Alternatively, the observed strain differences in social play levels with a novel playmate may be due to strain differences in responding to social isolation combined with exposure to a novel playmate. Long-Evans show lower social play levels with a novel playmate than Sprague-Dawley and Wistar after 48h isolation. It could be that Long-Evans experience social isolation as more stressful. In support, Long-Evans show higher hypothalamic-pituitary-adrenal axis (HPA) activity in response to acute stressors compared to Sprague-Dawley and Wistar (Sanchís-Ollé et al., 2021). Even under baseline conditions, Long-Evans rats have higher HPA axis activity than Wistar as shown by significantly higher peripheral plasma concentrations of ACTH and corticosterone (Tannahill et al., 1988). Whether these three rat strains differ in their hormonal and physiological responses to social isolation remains to be tested, but could be a first step toward understanding the neurobiological mechanisms underlying strain differences in social play levels when rats are exposed to a novel playmate.

### 2. Effects of playmate familiarity and social isolation length on social play tactics

#### Long-Evans and Sprague-Dawley show more role reversals with familiar playmates while Wistar show more role reversals with longer isolation time

Social play revolves around competition for species-typical bodily targets (Ham et al., 2024b), such as the nape of the neck in rats (Pellis and Pellis, 1987). For play bouts to be sustained, there must be some degree of reciprocity or turn taking (Palagi et al., 2016a; Pellis et al., 2024). In rats, play typically follows the ‘50:50’ rule (Altmann, 1962), with each rat in the pair acting in both the play offensive and play defensive roles equally (Palagi et al., 2016b). Indeed, rats will even engage in self-handicapping behaviors in order to maintain this even split in roles (Foroud and Pellis, 2003; Pellis et al., 2005). Here, we found that around 45-50% of playful attacks launched by both Long-Evans and Sprague-Dawley rats resulted in role reversals when exposed to a familiar playmate. This number dropped to around 20-30% when exposed to a novel playmate. In contrast, Wistar rats engaged in similar, albeit lower (10-25%) levels of role reversals regardless of playmate familiarity. Isolation time of the experimental rats influenced the proportion of role reversals in Sprague-Dawley and Wistar playmates with 48h versus 2h isolation resulting in fewer role reversals in Sprague-Dawley but more role reversals in Wistar. Role reversals reflect cooperation and reciprocity between playmates and are used to assess the quality of social play. For example, a lower proportion of role reversals was associated with impaired development of the prefrontal cortex and executive functions (Ham et al., 2024a; Schneider et al., 2016; Stark et al., 2022). Thus, our findings show that the quality of social play may be higher in Long-Evans with exposure to familiar playmates, in Sprague-Dawley with exposure to familiar playmates or upon short isolation, and in Wistar upon longer isolation.

#### Sprague-Dawley and Wistar defend more playful attacks than Long-Evans and do so by showing opposing defensive play tactics

Strain differences were observed for social play tactics, especially when exposed to novel playmates. Here, Sprague-Dawley and Wistar defended more playful attacks than Long-Evans. Sprague-Dawley did so by evading playful attacks, confirming previous findings (Himmler et al., 2014a, 2014b). In contrast, Wistar turned to face their attacker resulting in more complete rotations. These different defensive styles have opposing effects on the duration of body contact: evasion decreases body contact (the defender withdraws from the attacker) while facing defenses, especially followed by complete rotations, increases body contact as the rats wrestle one another. Notably, rats with neonatal damage to the motor cortex exhibited more evasions but fewer complete rotations than did control rats (Kamitakahara et al., 2007). Although the study used Long-Evans rats, their outcomes suggest that strain differences in social play tactics, as observed in our study, may be due to variations in the degree of motor cortex control over defensive play patterns, a possibility that could be tested in future studies.

All domesticated strains of rat originate from a common ancestor, a wild caught Norway rat (*Rattus norvegicus*), however, individual strains have diverse origins (Castle, 1947). Wistar rats come from the first domesticated lineage. From this Wistar strain, Long-Evans rats were developed by crossing several female Wistar rats with a single wild caught male rat (Castle, 1947; Lockard, 1968). Sprague-Dawley rats were derived by selective breeding of a line of Wistar rats (Castle, 1947; Krinke, 2000). Though both Long-Evans and Sprague-Dawley rats were first bred in the early 20th century, their variation in genetic background means that Sprague-Dawley rats are more closely related to Wistars than Long-Evans rats are. This could, in part, explain why most of the differences in defensive tactics, when partnered with a novel partner, are observed between the Long-Evans x Sprague-Dawley and Long-Evans x Wistar rats, and not between Sprague-Dawleys x Wistars.

#### Sex differences in social play tactics in Long-Evans, but not in Sprague-Dawley or Wistar

In rats, the proportion of attacks defended is similar between the sexes (Pellis, 2002), a result confirmed in all three rat strains in the present study. However, how males and females defend themselves was different in the Long-Evans strain. As Long-Evans rats age, weanlings switch from engaging in more partial rotations to more complete rotations once becoming juveniles (Pellis and Pellis, 1997). This is a change that both males and females exhibit. As males enter young adulthood, they switch back to preferring partial rotations over complete rotations. However, females continue to rotate fully when defending themselves as adults (Pellis, 2002). In our study, we found that female Long-Evans rats engage in more complete rotations than males do. This might reflect these sex-typical developmental differences, though at a younger age than what is typically observed, as all of the rats tested in this study were juveniles. The sex difference in complete rotations was not observed in Sprague-Dawley or Wistar rats suggesting that these sex-typical defensive tactics are specific for the Long-Evans strain.

### Conclusions

We showed that both playmate familiarity and social isolation length influenced social play behavior levels and tactics in juvenile rats, but did so differently for Long-Evans, Sprague-Dawley and Wistar. This resulted in robust strain differences with almost opposite playing preferences in Long-Evans versus Wistar, with Sprague-Dawley being the intermediate. Future research could aim to reveal the underlying neurobiological mechanisms linked to higher motivation to play with familiar playmates in Long-Evans and with novel playmates in Wistar. Given the common use of each of these rat strains in social play studies, our findings highlight the importance of taking into account the choice of strain in combination with external conditions like playmate familiarity and social isolation length. Finally, our findings are informative in suggesting that external conditions like playmate familiarity and social isolation length could influence social play levels and social play quality in typical and atypical children.

## ACKNOWLEDGEMENTS

This work was supported by the National Institutes of Health (R01MH125806 to AHV). JRH was supported by a PGS D grant from the Natural Science and Engineering Council of Canada (NSERC). We would like to thank Elie D.M Huez and Daniela Anderson for technical assistance.

## REFERENCES

Achterberg, E.J.M., Burke, C.J., Pellis, S.M., 2023. When the individual comes into play: The role of self and the partner in the dyadic play fighting of rats. Behav. Processes 212, 104933. 10.1016/j.beproc.2023.104933

Alessandri, S.M., 1992. Attention, play, and social behavior in ADHD preschoolers. J. Abnorm. Child Psychol. 20, 289–302. 10.1007/BF00916693

Altmann, S.A., 1962. Social behavior oxf anthropoid primates: Analysis of recent concepts., in: Roots of Behaviour. E.L. Bliss (Ed.), Hoeber, New York, pp. 277–285.

Argue, K.J., McCarthy, M.M., 2015. Characterization of juvenile play in rats: importance of sex of self and sex of partner. Biol. Sex Differ. 6, 16. 10.1186/s13293-015-0034-x

Barnett, L., 1990. Playfulness: Definition, design, and measurement, in: Play and Culture. pp. 319–336.

Beery, A.K., Shambaugh, K.L., 2021. Comparative Assessment of Familiarity/Novelty Preferences in Rodents. Front. Behav. Neurosci. 15, 648830. 10.3389/fnbeh.2021.648830

Bender, L., 1947. Childhood schizophrenia: Clinical study of one hundred schizophrenic children. Am. J. Orthopsychiatry 17, 40–56. 10.1111/j.1939-0025.1947.tb04975.x

Birke, L.I.A., Sadler, D., 1983. Progestin-induced changes in play behaviour of the prepubertal rat. Physiol. Behav. 30, 341–347. 10.1016/0031-9384(83)90136-1

Black, M., Freeman, B.J., Montgomery, J., 1975. Systematic observation of play behavior in autistic children. J. Autism Child. Schizophr. 5, 363–371. 10.1007/BF01540682

Bredewold, R., Nascimento, N.F., Ro, G.S., Cieslewski, S.E., Reppucci, C.J., Veenema, A.H., 2018. Involvement of dopamine, but not norepinephrine, in the sex-specific regulation of juvenile socially rewarding behavior by vasopressin. Neuropsychopharmacology 43, 2109–2117. 10.1038/s41386-018-0100-2

Bredewold, R., Schiavo, J.K., Van Der Hart, M., Verreij, M., Veenema, A.H., 2015. Dynamic changes in extracellular release of GABA and glutamate in the lateral septum during social play behavior in juvenile rats: Implications for sex-specific regulation of social play behavior. Neuroscience 307, 117–127. 10.1016/j.neuroscience.2015.08.052

Bredewold, R., Smith, C.J.W., Dumais, K.M., Veenema, A.H., 2014. Sex-specific modulation of juvenile social play behavior by vasopressin and oxytocin depends on social context. Front. Behav. Neurosci. 8. 10.3389/fnbeh.2014.00216

Calcagnetti, D.J., Schechter, M.D., 1992. Place conditioning reveals the rewarding aspect of social interaction in juvenile rats. Physiol. Behav. 51, 667–672. 10.1016/0031-9384(92)90101-7

Castle, W.E., 1947. The Domestication of the Rat. Proc. Natl. Acad. Sci. U. S. A. 33, 109–117. 10.1073/pnas.33.5.109

Chow, J.J., Beacher, N.J., Chabot, J.M., Oke, M., Venniro, M., Lin, D.-T., Shaham, Y., 2022. Characterization of operant social interaction in rats: effects of access duration, effort, peer familiarity, housing conditions, and choice between social interaction vs. food or remifentanil. Psychopharmacology (Berl.) 239, 2093–2108. 10.1007/s00213-022-06064-1

Darwin, C., Kebler, L., Joseph Meredith Toner Collection, 1871. The descent of man, : and selection in relation to sex. J. Murray, London.

Foroud, A., Pellis, S.M., 2003. The development of “roughness” in the play fighting of rats: A Laban Movement Analysis perspective. Dev. Psychobiol. 42, 35–43. 10.1002/dev.10088

Fujihara, K., Sato, T., Higeta, K., Miyasaka, Y., Mashimo, T., Yanagawa, Y., 2021. Behavioral Consequences of a Combination of Gad1 Haplodeficiency and Adolescent Exposure to an NMDA Receptor Antagonist in Long-Evans Rats. Front. Pharmacol. 12, 646088. 10.3389/fphar.2021.646088

Ginsburg, K.R., and the Committee on Communications, and the Committee on Psychosocial Aspects of Child and Family Health, 2007. The Importance of Play in Promoting Healthy Child Development and Maintaining Strong Parent-Child Bonds. Pediatrics 119, 182– 191. 10.1542/peds.2006-2697

Götz, F., Tönjes, R., Maywald, J., Dörner, G., 1991. Short- and Long-term Effects of a Dopamine Agonist (Lisuride) on Sex-specific Behavioural Patterns in Rats. Exp. Clin. Endocrinol. Diabetes 98, 111–121. 10.1055/s-0029-1211107

Hackenberg, T.D., Vanderhooft, L., Huang, J., Wagar, M., Alexander, J., Tan, L., 2021. Social preference in rats. J. Exp. Anal. Behav. 115, 634–649. 10.1002/jeab.686

Ham, J., Pellis, S., 2023. The Goldilocks Principle: Balancing Familiarity and Novelty in the Selection of Play Partners in Groups of Juvenile Male Rats. Anim. Behav. Cogn. 10, 304–328. 10.26451/abc.10.04.02.2023

Ham, J.R., Pellis, S.M., 2024. Play partner preferences among groups of unfamiliar juvenile male rats. Sci. Rep. 14, 16056. 10.1038/s41598-024-66988-w

Ham, J.R., Pellis, S.M., Pellis, V.C., 2024b. Oppositions, joints, and targets: the attractors that are the glue of social interactions. Front. Behav. Neurosci. 18, 1451283. 10.3389/fnbeh.2024.1451283

Ham, J.R., Szabo, M., Annor-Bediako, J., Stark, R.A., Iwaniuk, A.N., Pellis, S.M., 2024a. Quality not quantity: Deficient juvenile play experiences lead to altered medial prefrontal cortex neurons and sociocognitive skill deficits. Dev. Psychobiol. 66, e22456. 10.1002/dev.22456

Hawkins, P., Golledge, H.D.R., 2018. The 9 to 5 Rodent − Time for Change? Scientific and animal welfare implications of circadian and light effects on laboratory mice and rats. J. Neurosci. Methods 300, 20–25. 10.1016/j.jneumeth.2017.05.014

Helgeland, M.I., Torgersen, S., 2005. Stability and prediction of schizophrenia from adolescence to adulthood. Eur. Child Adolesc. Psychiatry 14, 83–94. 10.1007/s00787-005-0436-0

Himmler, B.T., Pellis, V.C., Pellis, S.M., 2013b. Peering into the Dynamics of Social Interactions: Measuring Play Fighting in Rats. J. Vis. Exp. 4288. 10.3791/4288

Himmler, B.T., Stryjek, R., Modlinska, K., Derksen, S.M., Pisula, W., Pellis, S.M., 2013a. How domestication modulates play behavior: A comparative analysis between wild rats and a laboratory strain of Rattus norvegicus. J. Comp. Psychol. 127, 453–464. 10.1037/a0032187

Himmler, S.M., Himmler, B.T., Pellis, V.C., Pellis, S.M., 2016. Play, variation in play and the development of socially competent rats. Behaviour 153, 1103–1137. 10.1163/1568539X-00003307

Himmler, S.M., Lewis, J.M., Pellis, S.M., 2014a. The Development of Strain Typical Defensive Patterns in the Play Fighting of Laboratory Rats. Int. J. Comp. Psychol. 27. 10.46867/ijcp.2014.27.03.09

Himmler, S.M., Modlinska, K., Stryjek, R., Himmler, B.T., Pisula, W., Pellis, S.M., 2014b. Domestication and diversification: A comparative analysis of the play fighting of the Brown Norway, Sprague-Dawley, and Wistar laboratory strains of (Rattus norvegicus). J. Comp. Psychol. 128, 318–327. 10.1037/a0036104

Hughes, E.M., Calcagno, P., Clarke, M., Sanchez, C., Smith, K., Kelly, J.P., Finn, D.P., Roche, M., 2020a. Prenatal exposure to valproic acid reduces social responses and alters mRNA levels of opioid receptor and pre-pro-peptide in discrete brain regions of adolescent and adult male rats. Brain Res. 1732, 146675. 10.1016/j.brainres.2020.146675

Hughes, E.M., Thornton, A.M., Kerr, D.M., Smith, K., Sanchez, C., Kelly, J.P., Finn, D.P., Roche, M., 2020b. Kappa Opioid Receptor-mediated Modulation of Social Responding in Adolescent Rats and in Rats Prenatally Exposed to Valproic Acid. Neuroscience 444, 9– 18. 10.1016/j.neuroscience.2020.07.055

Ikemoto, S., Panksepp, J., 1992. The effects of early social isolation on the motivation for social play in juvenile rats. Dev. Psychobiol. 25, 261–274. 10.1002/dev.420250404

Jessen, H.M., Kolodkin, M.H., Bychowski, M.E., Auger, C.J., Auger, A.P., 2010. The Nuclear Receptor Corepressor Has Organizational Effects within the Developing Amygdala on Juvenile Social Play and Anxiety-Like Behavior. Endocrinology 151, 1212–1220. 10.1210/en.2009-0594

Jones, P., 1994. Child developmental risk factors for adult schizophrenia in the British 1946 birth cohort. The Lancet 344, 1398–1402. 10.1016/S0140-6736(94)90569-X

Jones, P., Murray, R., Rodgers, B., 1995. Childhood Risk Factors for Adult Schizophrenia in a General Population Birth Cohort at Age 43 Years, in: Mednick, S.A., Hollister, J.M. (Eds.), Neural Development and Schizophrenia. Springer US, Boston, MA, pp. 151–176. 10.1007/978-1-4615-1955-3_10

Jordan, R., 2003. Social Play and Autistic Spectrum Disorders: A Perspective on Theory, Implications and Educational Approaches. Autism 7, 347–360. 10.1177/1362361303007004002

Kamitakahara, H., Monfils, M.-H., Forgie, M.L., Kolb, B., Pellis, S.M., 2007. The modulation of play fighting in rats: Role of the motor cortex. Behav. Neurosci. 121, 164–176. 10.1037/0735-7044.121.1.164

Kisko, T.M., Braun, M.D., Michels, S., Witt, S.H., Rietschel, M., Culmsee, C., Schwarting, R.K.W., Wöhr, M., 2018. *Cacna1c* haploinsufficiency leads to pro-social 50-kHz ultrasonic communication deficits in rats. Dis. Model. Mech. dmm.034116. 10.1242/dmm.034116

Krinke, G.J., 2000. The Laboratory Rat, 1st ed. Academic Press, San Diego, CA.

Ku, K.M., Weir, R.K., Silverman, J.L., Berman, R.F., Bauman, M.D., 2016. Behavioral Phenotyping of Juvenile Long-Evans and Sprague-Dawley Rats: Implications for Preclinical Models of Autism Spectrum Disorders. PLOS ONE 11, e0158150. 10.1371/journal.pone.0158150

Kurian, J.R., Bychowski, M.E., Forbes-Lorman, R.M., Auger, C.J., Auger, A.P., 2008. Mecp2 Organizes Juvenile Social Behavior in a Sex-Specific Manner. J. Neurosci. 28, 7137– 7142. 10.1523/JNEUROSCI.1345-08.2008

Lee, J.D.A., Reppucci, C.J., Huez, E.D.M., Bredewold, R., Veenema, A.H., 2024. Sex differences in the structure and function of the vasopressin system in the ventral pallidum are associated with the sex-specific regulation of social play behavior in juvenile rats. Horm. Behav. 163, 105563. 10.1016/j.yhbeh.2024.105563

Livia Terranova, M., Cirulli, F., Laviola, G., 1999. Behavioral and hormonal effects of partner familiarity in periadolescent rat pairs upon novelty exposure. Psychoneuroendocrinology 24, 639–656. 10.1016/S0306-4530(99)00019-0

Ljubetic, M., Maglica, T., Vukadin, Ž., 2020. BCES Conference Book 18.

Lockard, R.B., 1968. The albino rat: A defensible choice or a bad habit? Am. Psychol. 23, 734– 742. 10.1037/h0026726

Lukas, M., Wöhr, M., 2015. Endogenous vasopressin, innate anxiety, and the emission of pro-social 50-kHz ultrasonic vocalizations during social play behavior in juvenile rats. Psychoneuroendocrinology 56, 35–44. 10.1016/j.psyneuen.2015.03.005

Lundberg, S., Martinsson, M., Nylander, I., Roman, E., 2017. Altered corticosterone levels and social play behavior after prolonged maternal separation in adolescent male but not female Wistar rats. Horm. Behav. 87, 137–144. 10.1016/j.yhbeh.2016.11.016

Manduca, A., Campolongo, P., Palmery, M., Vanderschuren, L.J.M.J., Cuomo, V., Trezza, V., 2014. Social play behavior, ultrasonic vocalizations and their modulation by morphine and amphetamine in Wistar and Sprague-Dawley rats. Psychopharmacology (Berl.) 231, 1661–1673. 10.1007/s00213-013-3337-9

Marquardt, A.E., VanRyzin, J.W., Fuquen, R.W., McCarthy, M.M., 2023. Social play experience in juvenile rats is indispensable for appropriate socio-sexual behavior in adulthood in males but not females. Front. Behav. Neurosci. 16, 1076765. 10.3389/fnbeh.2022.1076765

Meaney, M.J., Dodge, A.M., William, B.W., 1981. Sex-Dependent Effects of Amygdaloid Lesions on the Social Play of Prepubertal Rats I. Meaney, M.J., McEwen, B.S., 1986. Testosterone implants into the amygdala during the neonatal period masculinize the social play of juvenile female rats. Brain Res. 398, 324– 328. 10.1016/0006-8993(86)91492-7

Meaney, M.J., Stewart, J., 1981. A descriptive study of social development in the rat (Rattus norvegicus). Anim. Behav. 29, 34–45. 10.1016/S0003-3472(81)80149-2

Meaney, M.J., Stewart, J., Poulin, P., McEwen, B.S., 1983. Sexual Differentiation of Social Play in Rat Pups Is Mediated by the Neonatal Androgen-Receptor System. Neuroendocrinology 37, 85–90. 10.1159/000123524

Mitchell, D., 1976. Experiments on neophobia in wild and laboratory rats: A reevaluation. J. Comp. Physiol. Psychol. 90, 190–197. 10.1037/h0077196

Moller, P., Husby, R., 2000. The Initial Prodrome in Schizophrenia: Searching for Naturalistic Core Dimensions of Experience and Behavior. Schizophr. Bull. 26, 217–232. 10.1093/oxfordjournals.schbul.a033442

Niesink, R.J.M., Van Ree, J.M., 1989. Involvement of opioid and dopaminergic systems in isolation-induced pinning and social grooming of young rats. Neuropharmacology 28, 411–418. 10.1016/0028-3908(89)90038-5

Nijhof, S.L., Vinkers, C.H., Van Geelen, S.M., Duijff, S.N., Achterberg, E.J.M., Van Der Net, J., Veltkamp, R.C., Grootenhuis, M.A., Van De Putte, E.M., Hillegers, M.H.J., Van Der Brug, A.W., Wierenga, C.J., Benders, M.J.N.L., Engels, R.C.M.E., Van Der Ent, C.K., Vanderschuren, L.J.M.J., Lesscher, H.M.B., 2018. Healthy play, better coping: The importance of play for the development of children in health and disease. Neurosci. Biobehav. Rev. 95, 421–429. 10.1016/j.neubiorev.2018.09.024

Northcutt, K.V., Nguyen, J.M.K., 2014. Female juvenile play elicits Fos expression in dopaminergic neurons of the VTA. Behav. Neurosci. 128, 178–186. 10.1037/a0035964

Northcutt, K.V., Nwankwo, V.C., 2018. Sex differences in juvenile play behavior differ among rat strains. Dev. Psychobiol. 60, 903–912. 10.1002/dev.21760

Olesen, K.M., Jessen, H.M., Auger, C.J., Auger, A.P., 2005. Dopaminergic Activation of Estrogen Receptors in Neonatal Brain Alters Progestin Receptor Expression and Juvenile Social Play Behavior. Endocrinology 146, 3705–3712. 10.1210/en.2005-0498

Olioff, M., Stewart, J., 1978. Sex differences in the play behavior of prepubescent rats. Physiol. Behav. 20, 113–115. 10.1016/0031-9384(78)90060-4

Palagi, E., Burghardt, G.M., Smuts, B., Cordoni, G., Dall’Olio, S., Fouts, H.N., Řeháková-Petrů, M., Siviy, S.M., Pellis, S.M., 2016b. Rough-and-tumble play as a window on animal communication. Biol. Rev. 91, 311–327. 10.1111/brv.12172

Palagi, E., Cordoni, G., Demuru, E., Bekoff, M., 2016a. Fair play and its connection with social tolerance, reciprocity and the ethology of peace. Behaviour 153, 1195–1216. 10.1163/1568539X-00003336

Panksepp, J., 2007. Can PLAY Diminish ADHD and Facilitate the Construction of the Social Brain? J. Can. Acad. Child Adolesc. Psychiatry 16, 57–66.

Panksepp, J., 1998. Affective Neuroscience: The Foundations of Human and Animal Emotions, online. ed. Oxford Academic, New York, NY.

Panksepp, J., 1981. The ontogeny of play in rats. Dev. Psychobiol. 14, 327–332. 10.1002/dev.420140405

Panksepp, J., Beatty, W.W., 1980. Social deprivation and play in rats. Behav. Neural Biol. 30, 197–206. 10.1016/S0163-1047(80)91077-8

Parent, C.I., Meaney, M.J., 2008. The influence of natural variations in maternal care on play fighting in the rat. Dev. Psychobiol. 50, 767–776. 10.1002/dev.20342

Paul, M.J., Terranova, J.I., Probst, C.K., Murray, E.K., Ismail, N.I., De Vries, G.J., 2014. Sexually dimorphic role for vasopressin in the development of social play. Front. Behav. Neurosci. 8. 10.3389/fnbeh.2014.00058

Pellis, S., Pellis, V., 2013. The Playful Brain: Venturing to the Limits of Neuroscience, ebook. ed. A Oneworld Book.

Pellis, S.M., 2002. Sex Differences in Play Fighting Revisited: Traditional and Nontraditional Mechanisms of Sexual Differentiation in Rats. Arch Sex Behav 31, 17–26. 10.1023/a:1014070916047

Pellis, S.M., Pellis, V.C., 1997. The prejuvenile onset of play fighting in laboratory rats (Rattus norvegicus). Dev. Psychobiol. 31, 193–205. 10.1002/(SICI)1098-2302(199711)31:3<193::AID-DEV4>3.0.CO;2-N

Pellis, S.M., Pellis, V.C., 1987. Play-fighting differs from serious fighting in both target of attack and tactics of fighting in the laboratory ratRattus norvegicus. Aggress. Behav. 13, 227– 242. 10.1002/1098-2337(1987)13:4<227::AID-AB2480130406>3.0.CO;2-C

Pellis, S.M., Pellis, V.C., Foroud, A., 2005. Play Fighting: Aggression, Affiliation, and the Development of Nuanced Social Skills. The Guilford Press, pp. 47–62.

Pellis, S.M., Pellis, V.C., Ham, J.R., 2024. Play fighting revisited: its design features and how they shape our understanding of its mechanisms and functions. Front. Ethol. 3, 1362052. 10.3389/fetho.2024.1362052

Pellis, S.M., Pellis, V.C., Ham, J.R., Achterberg, E.J.M., 2022. The rough-and-tumble play of rats as a natural behavior suitable for studying the social brain. Front. Behav. Neurosci. 16, 1033999. 10.3389/fnbeh.2022.1033999

Pellis, S.M., Pellis, V.C., Ham, J.R., Stark, R.A., 2023. Play fighting and the development of the social brain: The rat’s tale. Neurosci. Biobehav. Rev. 145, 105037. 10.1016/j.neubiorev.2023.105037

Pellis, S.M., Pellis, V.C., Himmler, B.T., 2014. How Play Makes for a More Adaptable Brain. A Comparative and Neural Perspective. Am. J. Play 7.

Pellis, S.M., Williams, L.A., Pellis, V.C., 2017. Adult-Juvenile Play Fighting in Rats: Insight into the Experiences that Facilitate the Development of Socio-Cognitive Skills. Int. J. Comp. Psychol. 30. 10.46867/ijcp.2017.30.00.14

Poole, T.B., Fish, J., 1975. An investigation of playful behaviour in *Rattus norvegicus* and *Mus musculus* (Mammalia). J. Zool. 175, 61–71. 10.1111/j.1469-7998.1975.tb01391.x

Potter, H.W., 1933. SCHIZOPHRENIA IN CHILDREN. Am. J. Psychiatry 89, 1253–1270. 10.1176/ajp.89.6.1253

Reinhart, C.J., McIntyre, D.C., Metz, G.A., Pellis, S.M., 2006. Play fighting between kindling-prone (fast) and kindling-resistant (slow) rats. J. Comp. Psychol. 120, 19–30. 10.1037/0735-7036.120.1.19

Sanchís-Ollé, M., Sánchez-Benito, L., Fuentes, S., Gagliano, H., Belda, X., Molina, P., Carrasco, J., Nadal, R., Armario, A., 2021. Male long-Evans rats: An outbred model of marked hypothalamic-pituitary-adrenal hyperactivity. Neurobiol. Stress 15, 100355. 10.1016/j.ynstr.2021.100355

Schiavi, S., Melancia, F., Carbone, E., Buzzelli, V., Manduca, A., Peinado, P.J., Zwergel, C., Mai, A., Campolongo, P., Vanderschuren, L.J.M.J., Trezza, V., 2020. Detrimental effects of the ‘bath salt’ methylenedioxypyrovalerone on social play behavior in male rats. Neuropsychopharmacology 45, 2012–2019. 10.1038/s41386-020-0729-5

Schneider, P., Bindila, L., Schmahl, C., Bohus, M., Meyer-Lindenberg, A., Lutz, B., Spanagel, R., Schneider, M., 2016. Adverse Social Experiences in Adolescent Rats Result in Enduring Effects on Social Competence, Pain Sensitivity and Endocannabinoid Signaling. Front. Behav. Neurosci. 10. 10.3389/fnbeh.2016.00203

Siviy, S.M., Eck, S.R., McDowell, L.S., Soroka, J., 2017. Effects of cross-fostering on play and anxiety in juvenile Fischer 344 and Lewis rats. Physiol. Behav. 169, 147–154. 10.1016/j.physbeh.2016.11.035

Siviy, S.M., Panksepp, J., 2011. In search of the neurobiological substrates for social playfulness in mammalian brains. Neurosci. Biobehav. Rev. 35, 1821–1830. 10.1016/j.neubiorev.2011.03.006

Smith, C.J.W., Wilkins, K.B., Li, S., Tulimieri, M.T., Veenema, A.H., 2018. Nucleus accumbens mu opioid receptors regulate context-specific social preferences in the juvenile rat. Psychoneuroendocrinology 89, 59–68. 10.1016/j.psyneuen.2017.12.017

Smith, C.J.W., Wilkins, K.B., Mogavero, J.N., Veenema, A.H., 2015. Social Novelty Investigation in the Juvenile Rat: Modulation by the μ-Opioid System. J. Neuroendocrinol. 27, 752– 764. 10.1111/jne.12301

Spinka, M., Newberry, R.C., Bekoff, M., 2001. Mammalian Play: Training for the Unexpected. Q. Rev. Biol. 76, 141–168. 10.1086/393866

Stark, R.A., 2021. THE ROLE OF PLAY-DERIVED EXPERIENCES ON THE DEVELOPMENT OF THE MEDIAL PREFRONTAL CORTEX AND ADULT SOCIAL BEHAVIOR. Univ. Lethbridge Can. ProQuest Diss. Theses.

Stark, R.A., Brinkman, B., Gibb, R.L., Iwaniuk, A.N., Pellis, S.M., 2022. Atypical play experiences in the juvenile period has an impact on the development of the medial prefrontal cortex in both male and female rats. Behav. Brain Res. 10.1016/j.bbr.2022.114222

Stark, R.A.M., Ramkumar, R., Pellis, S.M., 2021. Deficient Play-Derived Experiences in Juvenile Long Evans Rats Reared with a Fischer 344 Partner: A Deficiency Shared by Both Sexes. Int. J. Comp. Psychol. 34. 10.46867/IJCP.2021.34.5592

Supekar, K., Kochalka, J., Schaer, M., Wakeman, H., Qin, S., Padmanabhan, A., Menon, V., 2018. Deficits in mesolimbic reward pathway underlie social interaction impairments in children with autism. Brain. 10.1093/brain/awy191

Takahashi, L.K., Lore, R.K., 1983. Play fighting and the development of agonistic behavior in male and female rats. Aggress. Behav. 9, 217–227. 10.1002/1098-2337(1983)9:3<217::AID-AB2480090303>3.0.CO;2-4

Tannahill, L.A., Dow, R.C., Fairhall, K.M., Robinson, I.C.A.F., Fink, G., 1988. Comparison of Adrenocorticotropin Control in Brattleboro, Long-Evans, and Wistar Rats. Neuroendocrinology 48, 650–657. 10.1159/000125077

Thor, D.H., Holloway, W.R., 1984. Developmental analyses of social play behavior in juvenile rats. Bull. Psychon. Soc. 22, 587–590. 10.3758/BF03333916

Trezza, V., Baarendse, P.J.J., Vanderschuren, L.J.M.J., 2010. The pleasures of play: pharmacological insights into social reward mechanisms. Trends Pharmacol. Sci. 31, 463–469. 10.1016/j.tips.2010.06.008

Trezza, V., Vanderschuren, L.J.M.J., 2008. Cannabinoid and opioid modulation of social play behavior in adolescent rats: Differential behavioral mechanisms. Eur. Neuropsychopharmacol. 18, 519–530. 10.1016/j.euroneuro.2008.03.001

Vanderschuren, L., Niesink, R.J.M., Spruijt, B.M., Van Ree, J.M., 1995b. Influence of environmental factors on social play behavior of juvenile rats. Physiol. Behav. 58, 119– 123. 10.1016/0031-9384(94)00385-I

Vanderschuren, L.J.M.J., Achterberg, E.J.M., Trezza, V., 2016. The neurobiology of social play and its rewarding value in rats. Neurosci. Biobehav. Rev. 70, 86–105. 10.1016/j.neubiorev.2016.07.025

Vanderschuren, L.J.M.J., Niesink, R.J.M., Van Pee, J.M., 1997. The neurobiology of social play behavior in rats. Neurosci. Biobehav. Rev. 21, 309–326. 10.1016/S0149-7634(96)00020-6

Vanderschuren, L.J.M.J., Spruijt, B.M., Van Ree, J.M., Niesink, R.J.M., 1995a. Effects of morphine on different aspects of social play in juvenile rats. Psychopharmacology (Berl.) 117, 225–231. 10.1007/BF02245191

VanRyzin, J.W., Marquardt, A.E., McCarthy, M.M., 2020. Developmental origins of sex differences in the neural circuitry of play. Int. J. Play 9, 58–75. 10.1080/21594937.2020.1723370

Veenema, A.H., Bredewold, R., De Vries, G.J., 2013. Sex-specific modulation of juvenile social play by vasopressin. Psychoneuroendocrinology 38, 2554–2561. 10.1016/j.psyneuen.2013.06.002

Veenema, A.H., Neumann, I.D., 2009. Maternal separation enhances offensive play-fighting, basal corticosterone and hypothalamic vasopressin mRNA expression in juvenile male rats. Psychoneuroendocrinology 34, 463–467. 10.1016/j.psyneuen.2008.10.017

Win-Shwe, T.-T., Kyi-Tha-Thu, C., Fujitani, Y., Tsukahara, S., Hirano, S., 2021. Perinatal Exposure to Diesel Exhaust-Origin Secondary Organic Aerosol Induces Autism-Like Behavior in Rats. Int. J. Mol. Sci. 22, 538. 10.3390/ijms22020538

Wolfberg, P.J., Schuler, A.L., 1993. Integrated play groups: A model for promoting the social and cognitive dimensions of play in children with autism. J. Autism Dev. Disord. 23, 467– 489. 10.1007/BF01046051

Yogman, M., Garner, A., Hutchinson, J., Hirsh-Pasek, K., Golinkoff, R.M., COMMITTEE ON PSYCHOSOCIAL ASPECTS OF CHILD AND FAMILY HEALTH, COUNCIL ON COMMUNICATIONS AND MEDIA, Baum, R., Gambon, T., Lavin, A., Mattson, G., Wissow, L., Hill, D.L., Ameenuddin, N., Chassiakos, Y. (Linda) R., Cross, C., Boyd, R., Mendelson, R., Moreno, M.A., MSEd, Radesky, J., Swanson, W.S., MBE, Hutchinson, J., Smith, J., 2018. The Power of Play: A Pediatric Role in Enhancing Development in Young Children. Pediatrics 142, e20182058. 10.1542/peds.2018-2058

Yuill, N., Strieth, S., Roake, C., Aspden, R., Todd, B., 2007. Brief Report: Designing a Playground for Children with Autistic Spectrum Disorders––Effects on Playful Peer Interactions. J. Autism Dev. Disord. 37, 1192–1196. 10.1007/s10803-006-0241-8

