## Supplemental materials for "To Play or Not to Play? Effects of Playmate Familiarity and Social Isolation on Social Play Engagement in Three Laboratory Rat Strains"

### SUPPLEMENTARY RESULTS

#### Condition comparisons per strain for other social behaviors and cage exploration

***Long-Evans***

There was a significant main effect of Isolation on the duration of cage exploration (Suppl. Table 1), with overall more cage exploration in the 48h isolation period.

There was a significant Playmate x Isolation effect on the duration of social investigation, duration of total social behaviors, and duration of cage exploration (Suppl. Table 1). Bonferroni post hoc tests (Suppl. Table 2) showed that Long-Evans rats in the 2h-Novel condition have longer durations of social investigation and total social behaviors compared to the 2h-Familiar and 48h-Novel conditions (Suppl. Fig. 1A, 1C). In addition, Long-Evans rats in the 48h-Familiar condition have a longer duration of total social behaviors compared to the 2h-Familiar and 48h-Novel conditions (Suppl. Fig. 1C).

Furthermore, Long-Evans rats in the 2h-Novel condition showed less cage exploration compared to the 2h-Familiar and 48h-Novel conditions (Suppl. Fig. 1D) while the 48h-Novel condition has a longer duration of cage exploration compared to the 48h-Familiar condition (Suppl. Fig. 1D). There were no significant main effects on allogrooming nor any significant main effect of Sex or interaction effects with Sex on any of these behaviors (Suppl. Table 1).

***Sprague-Dawley***

There were significant main effects of Isolation and Playmate x Isolation (Suppl. Table 3) as well as Sex, Sex x Playmate and Sex x Playmate x Isolation (Suppl. Table 10) on the duration of allogrooming. Bonferroni post hoc tests showed that Sprague-Dawley rats in the 2h-Novel condition have a longer duration of allogrooming compared to the 48h-Novel and 2h-Familiar conditions (Suppl. Table 4; Suppl. Fig. 1F). This was driven by females showing more allogrooming in the 2h-Novel condition compared to males and compared to both the 48h-Novel and 2h-Familiar conditions (Suppl. Table 11).

There was a significant Playmate x Isolation effect on the duration of total social behaviors (Suppl. Table 3) with longer duration of total social behaviors in the 2h-Novel and 48h-Familiar conditions compared to their counterparts (Suppl. Table 4; Suppl Fig. 1G). There was also a significant Sex x Isolation interaction on the duration of total social behaviors (Suppl. Table 10) but Bonferroni post hoc testing did not reveal any further significance.

Last, there were significant main effects of Isolation and Playmate x Isolation on the duration of cage exploration (Suppl. Table 3) with less cage exploration in the 2h-Novel condition compared to the 2h-Familiar and 48h-Novel conditions (Suppl. Table 4; Suppl Fig. 1H).

***Wistar***
There was a significant Stimulus x Isolation interaction on the duration of total social behaviors (Suppl. Table 5) with shorter duration of total social behaviors in the 2h Familiar condition versus the 2h Novel and 48h Familiar conditions (Suppl. Fig. 1K, Suppl. Table 6).
There was a significant main effect of Isolation on the duration of cage exploration (Suppl. Table 5), with more cage exploration in the 48h isolation versus 2h isolation.

Finally, there was a significant main effect of Sex on the duration of allogrooming (Suppl. Table 5), with more allogrooming in males versus females.

**Strain comparisons per condition for other social behaviors and cage exploration**

***2h-Familiar Condition***
There were significant strain differences on the durations of social investigation, total social behaviors, and cage exploration (Suppl. Table 7). Bonferroni post hoc tests revealed that Wistar showed more social investigation than Long-Evans ($p = 0.017$; Suppl. Fig. 2A) but less cage exploration than Long-Evans and Sprague-Dawley (both: $p < 0.001$; Suppl. Fig. 2D). Bonferroni post hoc testing did not reveal any further significance for the duration of total social behaviors. Finally, there were no significant strain differences on the duration of allogrooming.

***48h-Familiar Condition***
There were significant strain differences on the durations of social investigation, total social behaviors, and cage exploration (Suppl. Table 7). Bonferroni post hoc tests revealed that Wistar showed more social investigation than Sprague-Dawley ($p = 0.007$; Suppl. Fig. 2E), showed more total social behaviors than Long-Evans ($p = 0.004$) and Sprague-Dawley ($p < 0.001$; Suppl. Fig. 2G) and showed less cage exploration than Long-Evans ($p = 0.024$) and Sprague-Dawley ($p < 0.001$; Suppl. Fig. 2H). There were no significant strain differences on the duration of allogrooming.

***2h-Novel Condition***
There were no significant strain differences on the durations of social investigation, allogrooming, and cage exploration. There was a significant strain difference on the duration of total social behaviors (Suppl. Table 7) but Bonferroni post hoc testing did not reveal any further significance between the strains.

***48h-Novel Condition***
There were significant strain differences on the durations of social investigation, total social behaviors, and cage exploration (Suppl. Table 7). Bonferroni post hoc tests revealed that Wistar rats show more social investigation (Suppl. Fig. 2M) and total social behaviors (Suppl. Fig. 2O)

but less cage exploration (Supp. Fig. 2P) compared to both Long-Evans ($p = 0.001$ for social investigation; $p < 0.001$ for total social behaviors and cage exploration) and Sprague-Dawley ($p = 0.013$ for social investigation; $p < 0.001$ for total social behaviors and cage exploration). There were no significant strain differences on the duration of allogrooming.

### SUPPLEMENTAL TABLES

**Supplemental Table 1. Experimental Condition comparisons for other social behaviors and cage exploration in experimental Long-Evans rats.** Three-way ANOVA statistics and partial eta squared (η2) effect sizes for the effects of Playmate (Familiar, Novel), Isolation (2h, 48h), and Sex on behavior of experimental male and female juvenile Long-Evans rats in the social play test. Significant effects and their corresponding behaviors are indicated in **bold**.

| | Playmate | Isolation | Playmate x Isolation | Sex |
|---|---|---|---|---|
| **Social investigation (% Time)** | $F_{(1,39)} = 2.07$, $p = 0.158$ | $F_{(1,39)} = 1.56$, $p = 0.220$ | **$F_{(1,39)} = 13.4$ $p < 0.001$ $\eta_p^2 = 0.26$** | $F_{(1,39)} = 0.10$, $p = 0.758$ |
| Allogrooming (% Time) | $F_{(1,39)} = 0.03$, $p = 0.872$ | $F_{(1,39)} = 1.19$, $p = 0.282$ | $F_{(1,39)} = 3.51$, $p = 0.069$ | $F_{(1,39)} = 1.18$, $p = 0.284$ |
| **Total social behaviors (% Time)** | $F_{(1,39)} = 2.55$, $p = 0.118$ | $F_{(1,39)} = 1.41$, $p = 0.243$ | **$F_{(1,39)} = 26.3$, $p < 0.001$ $\eta_p^2 = 0.40$** | $F_{(1,39)} = 0.04$, $p = 0.836$ |
| **Cage exploration (% Time)** | $F_{(1,39)} = 0.02$, $p = 0.879$ | **$F_{(1,39)} = 12.1$, $p = 0.001$ $\eta_p^2 = 0.24$** | **$F_{(1,39)} = 31.6$, $p < 0.001$ $\eta_p^2 = 0.45$** | $F_{(1,39)} = 0.00$, $p = 0.989$ |

**Supplemental Table 2. Posthoc tests for experimental condition comparisons for other social behaviors and cage exploration in experimental Long-Evans rats.** Bonferroni post hoc statistics and partial eta squared (η2) effect sizes for the interaction effects of Playmate (Familiar, Novel) and Isolation (2h, 48h) on behaviors of experimental male and female juvenile Long-Evans rats in the social play test. Significant effects and their corresponding behaviors are indicated in **bold**.

| | 48-Hour Familiar vs | | 2-Hour Novel vs | |
|---|---|---|---|---|
| | 2-Hour Familiar | 48-Hour Novel | 2-Hour Familiar | 48-Hour Novel |
| **Social investigation (% Time)** | $F_{(1,43)} = 3.73$, $p = 0.060$ | $F_{(1,43)} = 2.91$, $p = 0.095$ | **$F_{(1,43)} = 13.1$, $p < 0.001$ $\eta_p^2 = 0.23$** | **$F_{(1,43)} = 11.4$, $p = 0.002$ $\eta_p^2 = 0.21$** |
| **Total social behaviors (% Time)** | **$F_{(1,43)} = 9.73$, $p = 0.003$ $\eta_p^2 = 0.18$** | **$F_{(1,43)} = 27.0$, $p < 0.001$ $\eta_p^2 = 0.39$** | **$F_{(1,43)} = 6.05$, $p = 0.018$ $\eta_p^2 = 0.12$** | **$F_{(1,43)} = 18.7$, $p < 0.001$ $\eta_p^2 = 0.30$** |
| **Cage exploration (% Time)** | $F_{(1,43)} = 1.82$, $p = 0.185$ | **$F_{(1,43)} = 18.0$, $p < 0.001$ $\eta_p^2 = 0.30$** | **$F_{(1,43)} = 11.8$, $p = 0.001$ $\eta_p^2 = 0.21$** | **$F_{(1,43)} = 35.4$, $p < 0.001$ $\eta_p^2 = 0.45$** |

**Supplemental Table 3. Experimental Condition comparisons for other social behaviors and cage exploration in experimental Sprague-Dawley rats.** Three-way ANOVA statistics and partial eta squared (η2) effect sizes for the effects of Playmate (Familiar, Novel), Isolation (2h, 48h), and Sex on behavior of experimental male and female juvenile Sprague-Dawley rats in the social play test. Significant effects and their corresponding behaviors are indicated in **bold**.

| | Playmate | Isolation | Playmate x Isolation | Sex |
|---|---|---|---|---|
| Social investigation (% Time) | $F_{(1,41)} = 1.65$, $p = 0.207$ | $F_{(1,41)} = 1.31$, $p = 0.259$ | $F_{(1,41)} = 0.86$, $p = 0.360$ | $F_{(1,41)} = 1.38$, $p = 0.248$ |
| **Allogrooming (% Time)** | $F_{(1,41)} = 3.67$, $p = 0.063$ | **$F_{(1,41)} = 5.65$, $p = 0.022$ $\eta_p^2 = 0.12$** | **$F_{(1,41)} = 6.95$, $p = 0.012$ $\eta_p^2 = 0.14$** | **$F_{(1,41)} = 6.19$, $p = 0.017$ $\eta_p^2 = 0.13$** |
| **Total social behaviors (% Time)** | $F_{(1,41)} = 0.99$, $p = 0.325$ | $F_{(1,41)} = 0.21$, $p = 0.649$ | **$F_{(1,41)} = 16.1$, $p < 0.001$ $\eta_p^2 = 0.28$** | $F_{(1,41)} = 2.47$, $p = 0.123$ |
| **Cage exploration (% Time)** | $F_{(1,41)} = 1.08$, $p = 0.306$ | **$F_{(1,41)} = 15.4$, $p < 0.001$ $\eta_p^2 = 0.27$** | **$F_{(1,41)} = 9.84$, $p = 0.003$ $\eta_p^2 = 0.19$** | $F_{(1,41)} = 0.00$, $p = 0.999$ |

**Supplemental Table 4. Posthoc tests for experimental condition comparisons for other social behaviors and cage exploration in experimental Sprague-Dawley rats.** Bonferroni post hoc statistics and partial eta squared (η2) effect sizes for the interaction effects of Playmate (Familiar, Novel) and Isolation (2h, 48h) on behavior of experimental male and female juvenile Sprague-Dawley rats in the social play test. Significant effects and their corresponding behaviors are indicated in **bold**.

| | 48-Hour Familiar vs | | 2-Hour Novel vs | |
|---|---|---|---|---|
| | 2-Hour Familiar | 48-Hour Novel | 2-Hour Familiar | 48-Hour Novel |
| **Allogrooming (% Time)** | $F_{(1,45)} = 0.38$, $p = 0.540$ | $F_{(1,45)} = 0.20$, $p = 0.655$ | **$F_{(1,45)} = 8.16$, $p = 0.006$ $\eta_p^2 = 0.15$** | **$F_{(1,45)} = 7.86$, $p = 0.007$ $\eta_p^2 = 0.15$** |
| **Total social behaviors (% Time)** | **$F_{(1,45)} = 4.71$, $p = 0.035$ $\eta_p^2 = 0.09$** | **$F_{(1,45)} = 5.32$, $p = 0.026$ $\eta_p^2 = 0.11$** | **$F_{(1,45)} = 8.49$, $p = 0.006$ $\eta_p^2 = 0.16$** | **$F_{(1,45)} = 9.33$, $p = 0.004$ $\eta_p^2 = 0.17$** |
| **Cage exploration (% Time)** | $F_{(1,45)} = 0.06$, $p = 0.804$ | $F_{(1,45)} = 2.11$, $p = 0.154$ | **$F_{(1,45)} = 10.6$, $p = 0.002$ $\eta_p^2 = 0.19$** | **$F_{(1,45)} = 24.1$, $p < 0.001$ $\eta_p^2 = 0.35$** |

**Supplemental Table 5. Experimental Condition comparisons for other social behaviors and cage exploration in experimental Wistar rats.** Three-way ANOVA statistics and partial eta squared (η2) effect sizes for the effects of Playmate (Familiar, Novel), Isolation (2h, 48h), and Sex on behavior of experimental male and female juvenile Wistar rats in the social play test. Significant effects are indicated in **bold**.

| | Playmate | Isolation | Playmate x Isolation | Sex |
|---|---|---|---|---|
| Social investigation (% Time) | $F_{(1,72)} = 1.14$, $p = 0.288$ | $F_{(1,72)} = 0.02$, $p = 0.894$ | $F_{(1,72)} = 1.07$, $p = 0.304$ | $F_{(1,72)} = 2.90$, $p = 0.093$ |
| **Allogrooming (% Time)** | $F_{(1,72)} = 0.48$, $p = 0.489$ | $F_{(1,72)} = 0.32$, $p = 0.571$ | $F_{(1,72)} = 0.31$, $p = 0.580$ | **$F_{(1,72)} = 4.53$, $p = 0.037$ $\eta_p^2 = 0.06$** |
| **Total social behaviors (% Time)** | $F_{(1,72)} = 1.48$, $p = 0.228$ | $F_{(1,72)} = 2.84$, $p = 0.096$ | **$F_{(1,72)} = 7.37$, $p = 0.008$ $\eta_p^2 = 0.09$** | $F_{(1,72)} = 3.01$, $p = 0.087$ |
| **Cage exploration (% Time)** | $F_{(1,72)} = 1.26$, $p = 0.265$ | **$F_{(1,72)} = 8.06$, $p = 0.006$ $\eta_p^2 = 0.10$** | $F_{(1,72)} = 0.73$, $p = 0.394$ | $F_{(1,72)} = 0.13$, $p = 0.723$ |

**Supplemental Table 6. Posthoc tests for experimental condition comparisons for other social behaviors and cage exploration in experimental Wistar rats.** Bonferroni post hoc statistics and partial eta squared (η2) effect sizes for the interaction effects of Playmate (Familiar, Novel) and Isolation (2h, 48h) on behavior of experimental male and female juvenile Wistar rats in the social play test. Significant effects are indicated in **bold**.

| | 48-Hour Familiar vs | | 2-Hour Novel vs | |
|---|---|---|---|---|
| | 2-Hour Familiar | 48-Hour Novel | 2-Hour Familiar | 48-Hour Novel |
| **Total social behaviors (% Time)** | **$F_{(1,76)} = 9.69$, $p = 0.003$ $\eta_p^2 = 0.11$** | $F_{(1,76)} = 1.13$, $p = 0.292$ | **$F_{(1,76)} = 7.74$, $p = 0.007$ $\eta_p^2 = 0.09$** | $F_{(1,76)} = 0.53$, $p = 0.468$ |

**Supplemental Table 7. Strain comparisons for other social behaviors and cage exploration of experimental rats in each of the four experimental conditions.** One-way ANOVA statistics and partial eta squared (η2) effect sizes for the effect of Strain on behavior of experimental male and female juvenile Long-Evans, Sprague-Dawley, and Wistar rats in the social play test. Significant effects are indicated in **bold**.

| | 2-Hour Familiar | 48-Hour Familiar | 2-Hour Novel | 48-Hour Novel |
|---|---|---|---|---|
| **Social investigation (% Time)** | **$F_{(2,37)} = 4.29$, $p = 0.021$ $\eta_p^2 = 0.19$** | **$F_{(2,52)} = 5.14$, $p = 0.009$ $\eta_p^2 = 0.16$** | $F_{(2,37)} = 1.23$, $p = 0.291$ | **$F_{(2,38)} = 9.34$, $p < 0.001$ $\eta_p^2 = 0.33$** |
| Allogrooming (% Time) | $F_{(2,37)} = 0.83$, $p = 0.446$ | $F_{(2,52)} = 0.91$, $p = 0.407$ | $F_{(2,37)} = 0.69$, $p = 0.509$ | $F_{(2,38)} = 2.71$, $p = 0.079$ |
| **Total social behaviors (% Time)** | **$F_{(2,37)} = 3.74$, $p = 0.033$ $\eta_p^2 = 0.17$** | **$F_{(2,52)} = 10.6$, $p < 0.001$ $\eta_p^2 = 0.29$** | **$F_{(2,37)} = 3.43$, $p = 0.043$ $\eta_p^2 = 0.16$** | **$F_{(2,38)} = 30.9$, $p < 0.001$ $\eta_p^2 = 0.62$** |
| **Cage exploration (% Time)** | **$F_{(2,37)} = 15.0$, $p < 0.001$ $\eta_p^2 = 0.45$** | **$F_{(2,52)} = 13.1$, $p < 0.001$ $\eta_p^2 = 0.34$** | $F_{(2,37)} = 1.40$, $p = 0.259$ | **$F_{(2,38)} = 21.8$, $p < 0.001$ $\eta_p^2 = 0.53$** |

**Supplemental Table 8. Playmate x Isolation x Sex effect for role reversals in playmate Long-Evans rats.** Three-way ANOVA statistics and partial eta squared (η2) effect sizes for Playmate (Familiar, Novel), Isolation (2h, 48h), and Sex on the proportion of role reversals of playmate male and female juvenile Long-Evans rats in the social play test. Significant effects and their corresponding behaviors are indicated in **bold**.

| | Playmate x Isolation x Sex |
|---|---|
| **Role reversals** | **$F_{(1,39)} = 7.39$, $p = 0.010$ $\eta_p^2 = 0.16$** |

**Supplemental Table 9. Posthoc tests for Playmate x Isolation x Sex effect for role reversals in playmate Long-Evans rats.** Bonferroni post hoc statistics and partial eta squared (η2) effect sizes for the interaction effect of Playmate (Familiar, Novel), Isolation (2h, 48h), and Sex on the proportion of role reversals of playmate male and female juvenile Long-Evans rats in the social play test. Significant effects are indicated in **bold**.

| Role reversals | | | | |
|---|---|---|---|---|
| | 48-Hour Familiar vs | | 2-Hour Novel vs | |
| | 2-Hour Familiar | 48-Hour Novel | 2-Hour Familiar | 48-Hour Novel |
| Females | $F_{(1,39)} = 0.32$, $p = 0.573$ | **$F_{(1,39)} = 6.50$, $p = 0.015$ $\eta_p^2 = 0.14$** | $F_{(1,39)} = 0.57$, $p = 0.457$ | **$F_{(1,39)} = 6.33$, $p = 0.016$ $\eta_p^2 = 0.14$** |
| Males | $F_{(1,39)} = 1.37$, $p = 0.249$ | $F_{(1,39)} = 0.05$, $p = 0.825$ | **$F_{(1,39)} = 4.70$, $p = 0.036$ $\eta_p^2 = 0.11$** | $F_{(1,39)} = 0.98$, $p = 0.328$ |

**Supplemental Table 10. Sex interaction effects for other social behaviors in experimental Sprague-Dawley rats.** Three-way ANOVA statistics and partial eta squared (η2) effect sizes for the effects of Sex interaction effects on other social behaviors of experimental male and female juvenile Sprague-Dawley rats in the social play test. Significant effects and their corresponding behaviors are indicated in **bold**.

| | Sex x Playmate | Sex x Isolation | Sex x Playmate x Isolation |
|---|---|---|---|
| **Allogrooming (% Time)** | **$F_{(1,41)} = 9.78$, $p = 0.003$ $\eta_p^2 = 0.19$** | $F_{(1,41)} = 1.23$, $p = 0.274$ | **$F_{(1,41)} = 11.5$, $p = 0.002$ $\eta_p^2 = 0.22$** |
| **Total social behaviors (% Time)** | $F_{(1,41)} = 1.60$, $p = 0.213$ | **$F_{(1,41)} = 7.57$, $p = 0.009$ $\eta_p^2 = 0.16$** | $F_{(1,41)} = 1.56$, $p = 0.219$ |

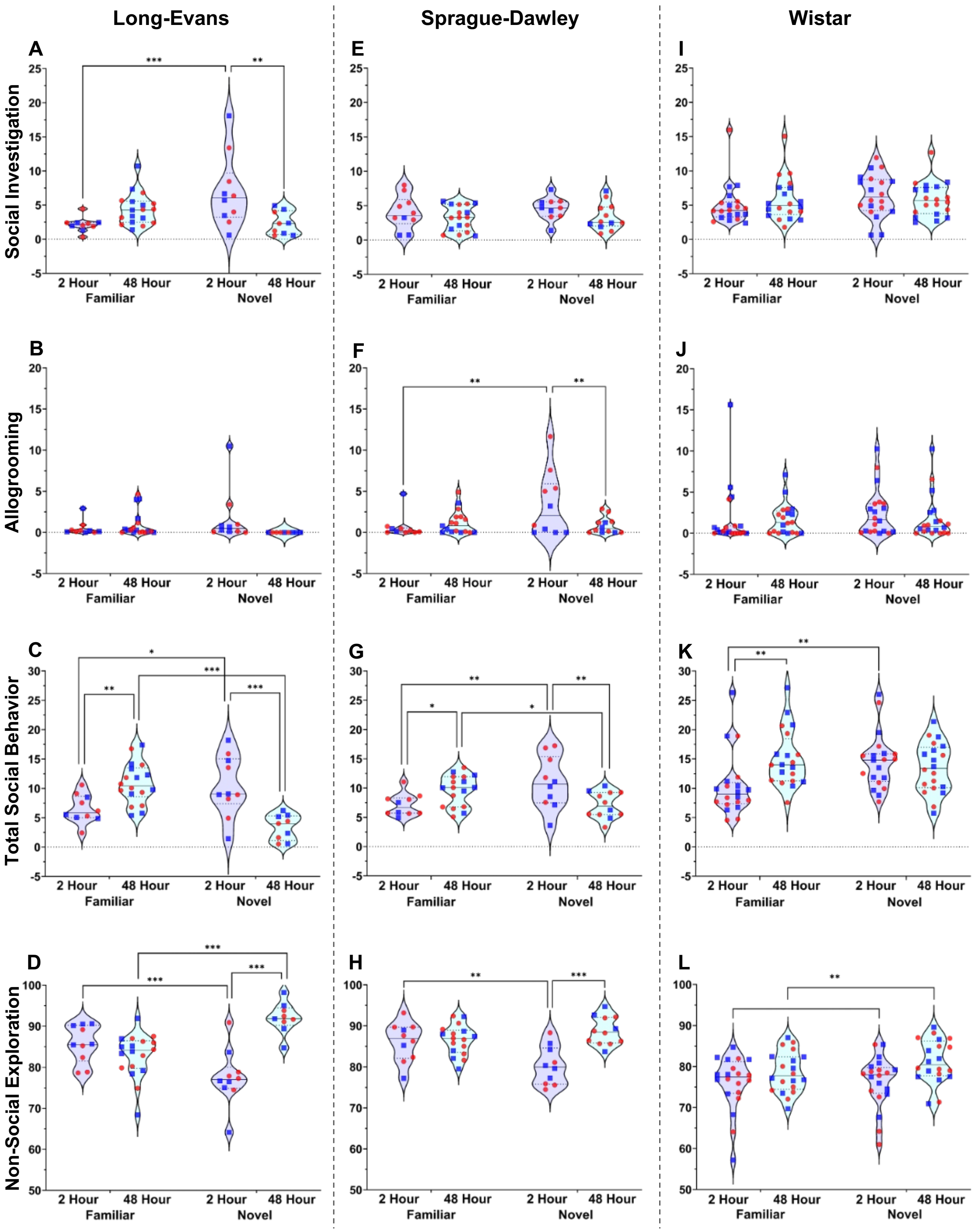


**Supplemental Fig. 1 | Experimental condition comparisons for other social and non-social behaviors per rat strain. A-D:** Long-Evans exposed to a novel playmate after a 2h isolation showed longer durations of social investigation (**A**) and total social behavior (**C**) but shorter duration of cage exploration

(**D**) compared to a familiar playmate and compared to 48h isolation. Moreover, Long-Evans exposed to a familiar playmate after 48h isolation showed longer duration of total social behavior (**C**) compared to 2h isolation and compared to a novel playmate while showing a shorter duration of exploration (**D**) compared to a novel playmate. No condition effects were found for allogrooming (**B)**. **E-H:** Sprague-Dawley did not show condition effects on social investigation duration (**E**). However, Sprague-Dawley exposed to a novel playmate after a 2h isolation showed longer durations of allogrooming (**F**) and total social behavior (**G**) but shorter duration of cage exploration (**H**) compared to a familiar playmate and compared to 48h isolation. Sprague-Dawley exposed to a familiar playmate after 48h isolation showed longer duration of total social behavior (**G**) compared to a novel playmate and compared to 2h isolation. **I-L:** Wistar did not show condition effects on social investigation (**I**) and allogrooming (**J**) durations. However, Wistar exposed to a familiar playmate after 2h isolation showed shorter duration of total social behavior (**K**) compared to a novel playmate and compared to 48h isolation, and longer duration of cage exploration (**L**) compared to a novel playmate. Moreover, Wistar exposed to a familiar playmate after 48h isolation showed shorter duration of cage exploration (**L**) compared to a novel playmate. The behaviors in the graphs are indicated as percentage of total time. The percentage duration of total social behavior is the summation of the percentage durations of social play, social investigation, and allogrooming; Males are represented by blue squares, females are represented by red circles; Three-way ANOVA analyses per strain to assess effects of Playmate, Isolation, and Sex, followed by Bonferroni post hoc tests to assess interaction effects; * $p < 0.05$, ** $p \leq 0.01$, *** $p \leq 0.001$.

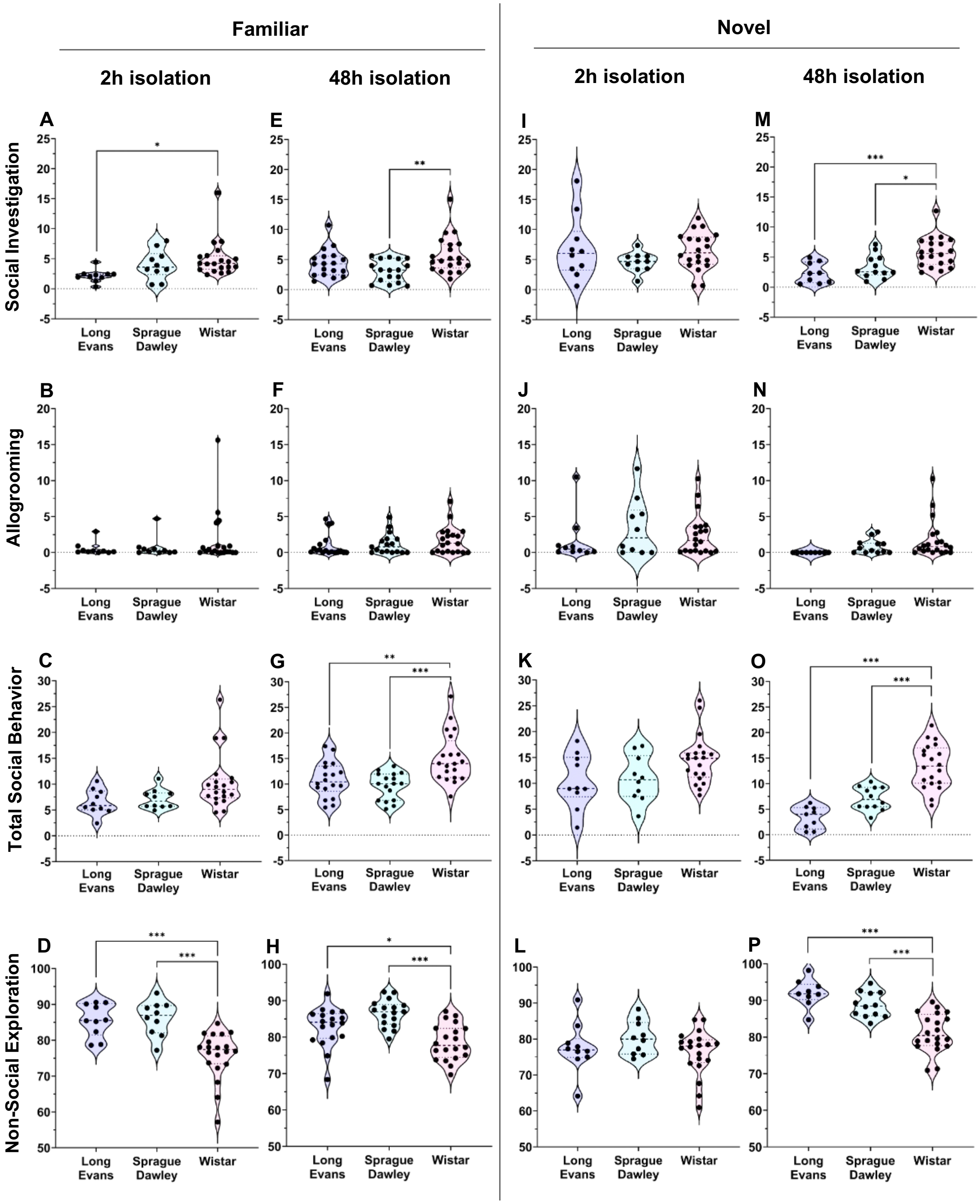


**Supplemental Fig. 2 | Rat strain comparisons for other social and non-social behaviors per experimental condition. A-D:** When rats were exposed to a familiar playmate after 2h isolation, Long-Evans showed less social investigation (**A**) but more cage exploration (**D**) than Wistar. Sprague Dawley also showed more cage exploration (**D**) than Wistar. No strain differences were found for durations of allogrooming (**B**) and total social behaviors (**C**). **E-H:** When rats were exposed to a familiar playmate after 48h isolation, Sprague-Dawley showed less social investigation (**E**) than Wistar. Long-Evans and

Sprague-Dawley showed less total social behavior (**G**) but more cage exploration (**H**) than Wistar. No strain differences were found for durations of allogrooming (**F**). **I-L:** No strain differences were found for durations of social investigation, allogrooming, total social behaviors and cage exploration when rats were exposed to a novel playmate after 2h isolation. **M-P:** When rats were exposed to a novel playmate after 48h isolation, Long-Evans and Sprague-Dawley showed less social investigation (**M**), less total social behavior (**O**), but more cage exploration (**P**) than Wistar. No strain differences were found for allogrooming (**N**). The behaviors in the graphs are indicated as percentage of total time. The percentage duration of total social behavior is the summation of the percentage durations of social play, social investigation, and allogrooming. One-way ANOVA analysis per experimental condition followed by Bonferroni post hoc tests to assess strain differences; * $p < 0.05$, ** $p \leq 0.01$, *** $p \leq 0.001$.
